# A role for circular code properties in translation

**DOI:** 10.1101/2020.09.28.317016

**Authors:** Simone Giannerini, Alberto Danielli, Diego Luis Gonzalez, Greta Goracci

## Abstract

Circular codes represent a form of coding allowing detection/correction of frameshift errors. Building on recent theoretical advances on circular codes, we provide evidence that protein coding sequences exhibit in-frame circular code marks, that are absent in introns and are intimately linked to the keto-amino transformation of codon bases. These properties strongly correlate with translation speed, codon influence and protein expression levels. Strikingly, circular code marks are absent at the beginning of coding sequences, but stably occur 40 codons after the initiator codon, hinting at the translation elongation process. Finally, we use the lens of circular codes to show that codon influence on translation correlates with the strong-weak dichotomy of the first two bases of the codon. The results provide promising universal tools for sequence indicators and sequence optimization for bioinformatics and biotechnological applications, and can shed light on the molecular mechanisms behind the decoding process.

## 1. Introduction

The genetic code is nearly universal across all living organisms. Its degeneracy, mapping 64 three-letter codons to 20 amino acids and three stop codons, is highly conserved. This conservation has evolved to minimize the effects of genetic mutations and translational decoding errors, thus providing optimal robustness in the flow of the genetic information Woese (1965). Moreover, the universal genetic code allows to tolerate arbitrary nucleotide sequences within protein-coding regions better than other possible codes. This property appears to reduce the deleterious effects of frame-shift translation errors, increasing the probability that a stop codon is encountered after a frame shift Itzkovitz & Alon (2007).

The degeneracy of the code enables to synthesize the same protein from a huge number of mRNA sequences encompassing synonymous codons. Growing evidence suggests that synonymous codons in coding sequences are not neutral with respect to the translation process, influencing the protein expression rates from bacteria to eukarya. It is generally accomplished that this codon bias contributes to the translation efficiency at the elongation step (Quax et al., 2015), which in turn may affect the stability of the translated mRNA Boël et al. (2016). As such, codon preferences have been intensively studied, both for improved protein production yields in biotechnological settings as well as for codon-optimized gene design in synthetic biology and genetic engineering projects Brule & Grayhack (2017).

The mechanistic effects of codon bias were initially attributed to inefficient translation of sets of rare codons Chen & Inouye (1994), implying the co-variance of codon usage frequency with the levels of matching tRNA pools and the deriving attenuation of translation elongation rates at infrequently used codons (Ikemura, 1981). The recent introduction of genome-wide ribosome profiling studies has questioned this simplistic view, since the net ribosome elongation rates are apparently relatively constant and marginally affected by rare codon frequency Ingolia (2014); Pop et al. (2014). On the other hand, several studies correlated the effect of codon bias either with the stability of secondary structures at the 5’ end of the mRNA Kudla et al. (2009); Bentele et al. (2013); Goodman et al. (2013), or with the intracistronic occurrence of Shine-Dalgarno-like sequences mimicking the ribosome binding site Li et al. (2012). Moreover, different types of codon bias have been described, including synonymous codon co-occurrence, allowing for rapid recycling of the exhaust tRNA in highly expressed genes Cannarrozzi et al. (2010), or non-synonymous codon pair bias, dependent on optimal interactions of tRNAs in the A and P sites of the ribo-some Demeshkina et al. (2012); Quax et al. (2015); Hanson & Coller (2018).

An elegant study engineered a *his* operon leader peptide gene reporter in *E. coli* to investigate the local effects of codon context on in vivo translation speed Chevance et al. (2014). Results demonstrated that the rate at which ribosomes translate individual synonymous codons varies considerably, and that the apparent speed at which a given codon is translated is influenced by flanking ones.

Recently, the codon influence on protein expression rates was assayed in greater depth, by integrating statistical analyses of large scale protein expression data sets with a systematic evaluation of local and global mRNA properties Gardin et al. (2014); Boël et al. (2016); Cambray et al. (2018). In particular, in Boël et al. (2016), a logistic regression model is used to build a codon-influence metric, validated by biochemical experiments, demonstrating that codon content is able to modulate the kinetic competition between translation elongation rates and mRNA stability. mRNA-folding effects generally prevail at the 5’ end of the coding sequence Kudla et al. (2009) and appear to be cumulatively weaker than codon bias effects Boël et al. (2016). Finally, it was shown that a major determinant of mRNA half-life and stability is the codon-optimized rate of translational elongation Presnyak et al. (2015).

Despite these advances, the theoretical principles behind the empirical effects of codon bias on translation efficiency remain poorly addressed. A possible correlative link between codon bias and reading frame maintenance was inferred from the statistical analysis of a large set of protein coding sequences in the three possible reading frames, resulting in the discovery that the set of most frequent in-frame codons formed a circular code Arquès & Michel (1996); Michel (2008, 2015). This observation revived the study of protein expression from the point of view of coding theory initiated by Crick with the introduction of comma-free codes Crick et al. (1957); Golomb et al. (1958). Recent developments on the theory of circular codes led to postulate the existence of a coding strategy underlying the process of reading frame maintenance Gonzalez et al. (2011); Fimmel et al. (2015a, b, 2016). Circular codes have been proposed as putative remnants of primeval comma-free codes Shepherd (1981); Dila et al. (2019a). The circular code found in Arquès & Michel (1996) belongs to a set of 216 codes possessing desirable properties (i.e. self-complementary, maximal, C^3^ circular codes, see Supplementary Information). In Fimmel et al. (2015a), it shown that such set can be partitioned into 27 equivalence classes conforming to a group theoretic framework characterized by 8 nucleotide transformations that are isomorphic to the symmetries of the square. Table 1 shows an example of an equivalence class formed by 8 circular codes linked by such transformations. It has been postulated that this mathematical structure could be correlated with the correct transmission of information and frame maintenance during translation Gonzalez et al. (2011); Michel (2012). Such premises encouraged us to investigate more thoroughly whether circular codes could provide a theoretical framework able to explain or predict the effects of codon bias on translation. Up to now, the key parameter, used to investigate the role of circular code properties on translation, has been represented by the **coverage** of a circular code over a specific sequence or organism. It is the cumulative codon usage of the set of codons belonging to that code:

**Table 1:**
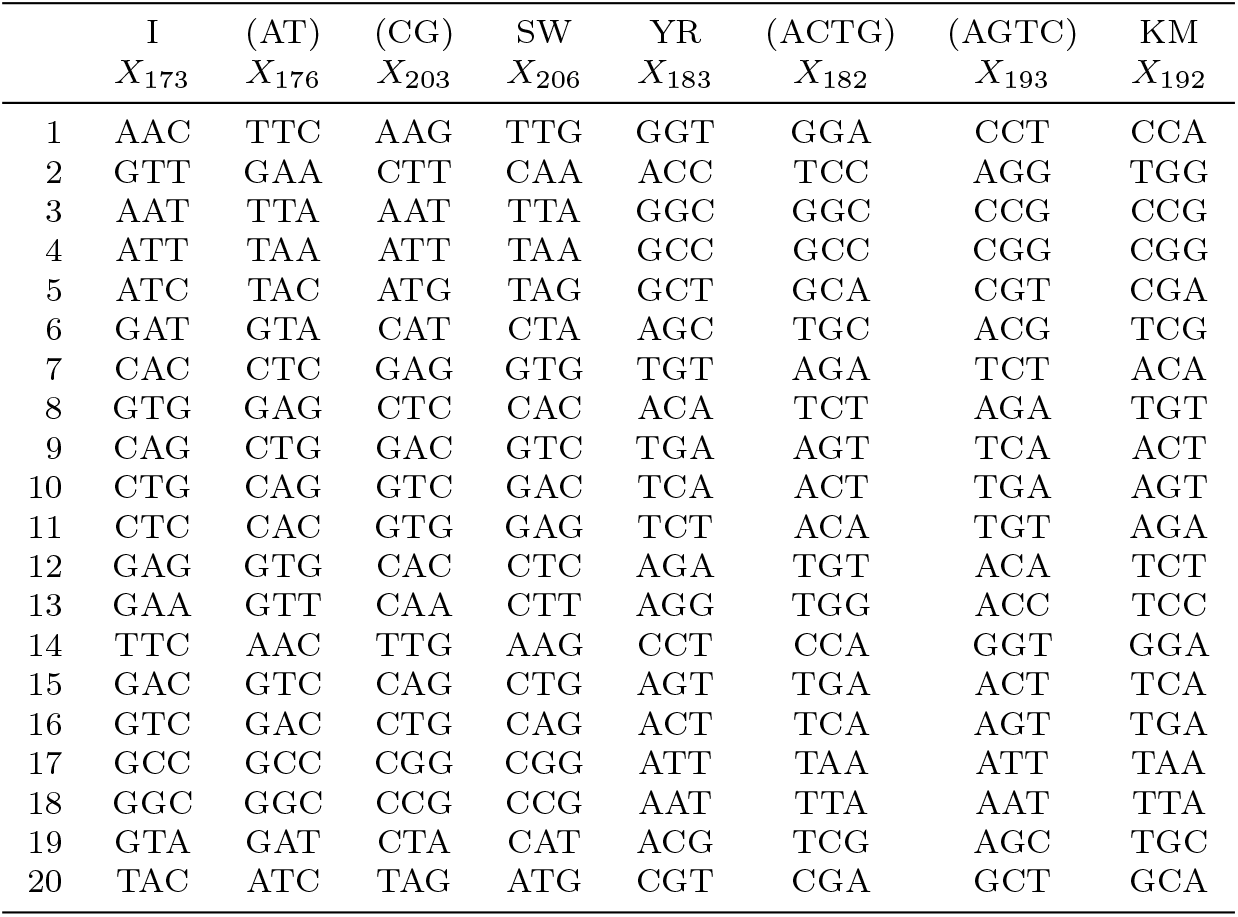
Equivalence class formed by eight circular codes. Each column represents one of the 216 circular codes. The codes are related through the group of transformations *D*_8_. For instance AAC ∈ *X*_173_ and KM(AAC) = CCA ∈ *X*_192_, see the Supplementary Information for details.

**Example 1.** Consider the sequence CAT CTG AAT GGA CTG and the two codes *X*_1_ = {CTG, AAT}, *X*_2_ = {GGA, TGT}. The coverage of *X*_1_ results 3/5 = 0.60, and that of *X*_2_ results 1/5 = 0.20.

Hence, the coverage of a code is the sum of the codon usages of its codons and can be seen as a measure of its “compliance” with the coding sequence, see also Gonzalez et al. (2011). For a rigourous mathematical definition of coverage and a brief description of circular codes theory see Supplementary Information.

In order to explore the relationship of circular codes with extant coding sequences, we set out to systematically compare the coverage of the 216 circular codes partitioned in 27 equivalence classes, with the codon usage of a large set of organisms.

## 2. Results and discussion

### 2.1. Circular codes’ coverage exhibits universal properties

We have analyzed the whole Codon Usage Database (https://www.kazusa.or.jp/codon/) to show the coverage (in percentage) for the 216 circular codes partitioned in 27 equivalence classes Fimmel et al. (2015a). As a paradigmatic example we present the results for 8 codes forming the equivalence class of Table 1 (the results for the remaining classes are reported in the Supplementary Information). The results are presented in Table 2. As expected, each code has a distinct degree of coverage reflecting taxon-specific codon usage. For instance, code *X*_173_ covers very well bacteria, i.e. the 46.4% of the codons of all bacterial genomes belong to code *X*_173_. In contrast, the coverage for plants is lower (39.7%). Such disparity is reflected in the absolute ranks shown in the middle panel: for bacteria, code *X*_173_ ranks 2^nd^ among the 216 codes whereas for plants it ranks 16^th^. This heterogeneity is evident also for the other 7 codes of the class for all the kingdoms. However, if we consider the ranks of these coverages inside the equivalence class (lower panel), then a neat taxon-independent ordering among the 8 codes emerges i.e. in this case, code *X*_173_ is always the best of its class, code *X*_176_ is always the second etc., irrespective of the species-specific codon usage. Surprisingly, this property holds for each of the 27 equivalence classes (see Table S5). Even more remarkably, the worst code within each class (code with the least coverage) invariably coincides with the chemical Keto-Amino transformation of the best one. In the example of Table 2, code *X*_173_ is always the best code and its Keto-Amino transformation KM(*X*_173_) = *X*_192_ is always the worst within the class. This establishes an important link between the codon usage and the Keto-Amino (KM) chemical transformation that will be discussed below. This property is not the trivial consequence of the fact that the more a set of codons is recurrent then, the less recurrent are codons that do not belong to the same set (see Supplementary Information, Section 2.1.1 where we set up a statistical test).

**Table 2:**
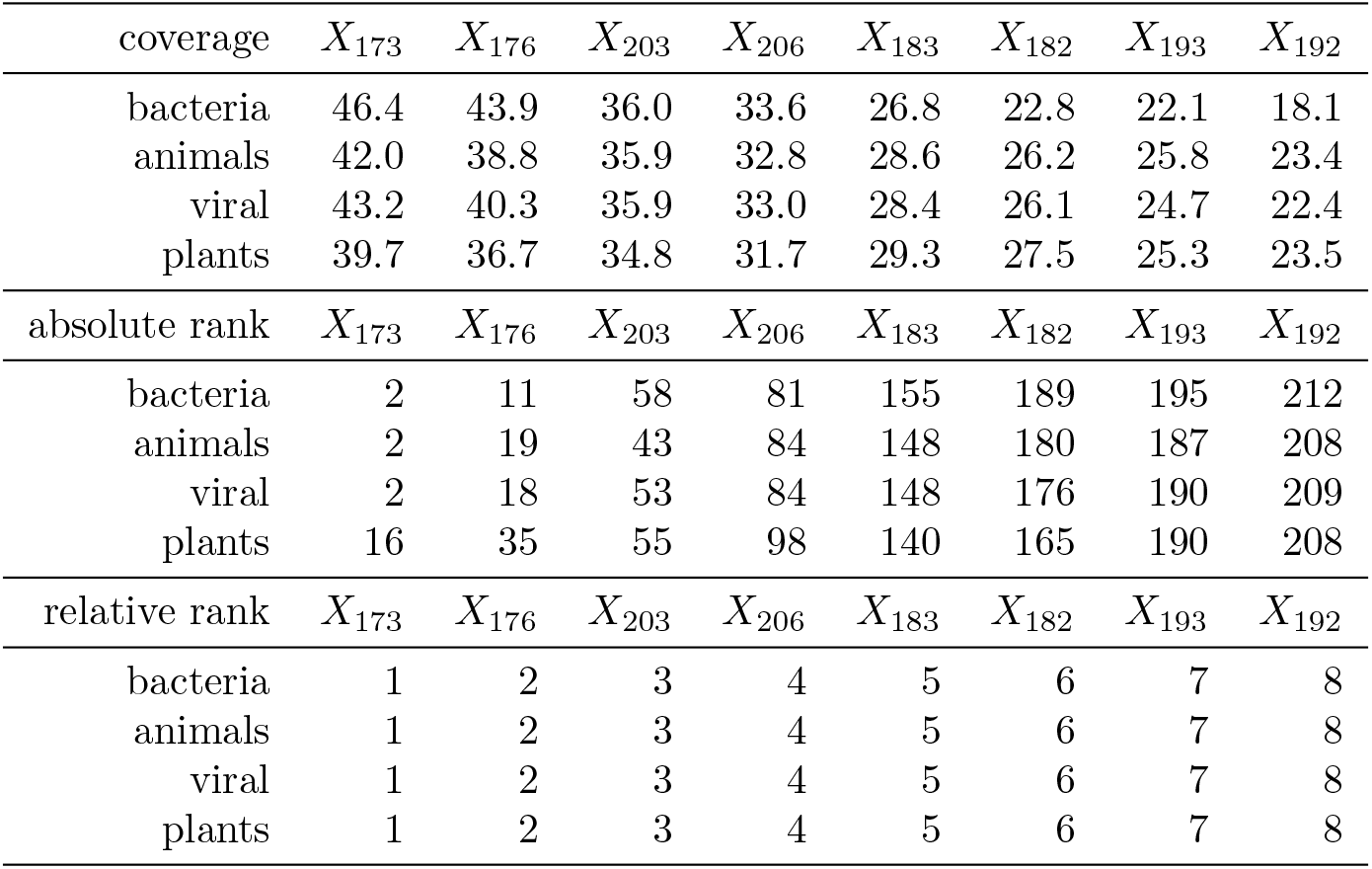
Coverage (upper panel), absolute ranks (mid panel) and relative ranks (lower panel) for the equivalence class of 8 circular codes presented in Table 1. The universality of the results is clear if we consider the ranks within classes: for instance the coverage of code *X*_173_ for bacteria is 46.4% (upper panel). It is not the highest coverage among the 216 codes, indeed it is the second (mid panel). However, it is always the highest within its class (lower panel). This universal behaviour holds for the whole set of 216 codes partitioned in 27 equivalence classes.

| coverage | $X_{173}$ | $X_{176}$ | $X_{203}$ | $X_{206}$ | $X_{183}$ | $X_{182}$ | $X_{193}$ | $X_{192}$ |
| --- | --- | --- | --- | --- | --- | --- | --- | --- |
| bacteria | 46.4 | 43.9 | 36.0 | 33.6 | 26.8 | 22.8 | 22.1 | 18.1 |
| animals | 42.0 | 38.8 | 35.9 | 32.8 | 28.6 | 26.2 | 25.8 | 23.4 |
| viral | 43.2 | 40.3 | 35.9 | 33.0 | 28.4 | 26.1 | 24.7 | 22.4 |
| plants | 39.7 | 36.7 | 34.8 | 31.7 | 29.3 | 27.5 | 25.3 | 23.5 |
| absolute rank | $X_{173}$ | $X_{176}$ | $X_{203}$ | $X_{206}$ | $X_{183}$ | $X_{182}$ | $X_{193}$ | $X_{192}$ |
| bacteria | 2 | 11 | 58 | 81 | 155 | 189 | 195 | 212 |
| animals | 2 | 19 | 43 | 84 | 148 | 180 | 187 | 208 |
| viral | 2 | 18 | 53 | 84 | 148 | 176 | 190 | 209 |
| plants | 16 | 35 | 55 | 98 | 140 | 165 | 190 | 208 |
| relative rank | $X_{173}$ | $X_{176}$ | $X_{203}$ | $X_{206}$ | $X_{183}$ | $X_{182}$ | $X_{193}$ | $X_{192}$ |
| bacteria | 1 | 2 | 3 | 4 | 5 | 6 | 7 | 8 |
| animals | 1 | 2 | 3 | 4 | 5 | 6 | 7 | 8 |
| viral | 1 | 2 | 3 | 4 | 5 | 6 | 7 | 8 |
| plants | 1 | 2 | 3 | 4 | 5 | 6 | 7 | 8 |

These results demonstrate that universal symmetry properties of coding sequences emerge when analyzed through the theoretical framework of circular codes, irrespectively of the species-specific codon-usage. Moreover, within each equivalence class, the Keto-Amino transformation of the code possessing the best coverage always leads to the worst covering code of the same class. Thus, a universal ordering structure, conserved across domains of life, emerges beyond the heterogeneity of species-specific codon usage.

### 2.2. Universal frame marks in coding sequences

The biological functions associated with circular code properties are basically unexplored. These properties may be explained as a fossilized memory of comma-free (self-synchronizable) coding in primeval forms of life Dila et al. (2019a), or tentatively associated with reading frame maintenance during protein synthesis Dila et al. (2019b). Thus, to explore whether the universal ranking property shown above, is valid also out of frame, we extended the analysis of the coverage of circular codes to the three reading frames of coding sequences for 25 well-annotated eukaryotic species (Table S6). The results are shown in the Supplementary Information, Table S7-S12. Remarkably, despite the variability of the codon usage among the different species, the ranking within each equivalence class is always preserved in the three frames. For example, for frame + 1, Tables S9-S10 (+2, Tables S11-S12), the first (second) circular permutation of the best codes has always the highest coverage, whereas their keto-amino transformation always leads to the worst covering codes within their equivalence class.

When ordered through the ranks, the coverage shows a strong linear scaling. This is shown in Figure 1 (left) that reports the boxplots of the coverage (percent) of the 8 circular codes of Table 1 over the in-frame coding sequences of the 25 eukaryotic genomes analysed. The same linear scaling is observed for the coverage of the first and second circularly permuted codes, over the same coding sequences read out-of-frame +1 (central panel) and +2 (right panel), respectively. Scaling laws are important in Information Theory (Wallace & Wallace, 1998) and dynamical system theory (Feigenbaum, 1988) and have also been associated to universal properties and long range correlations in DNA Cristadoro et al. (2018). Intriguingly, the structure uncovered in coding sequences is completely absent in introns (Table S13).

**Figure 1:**
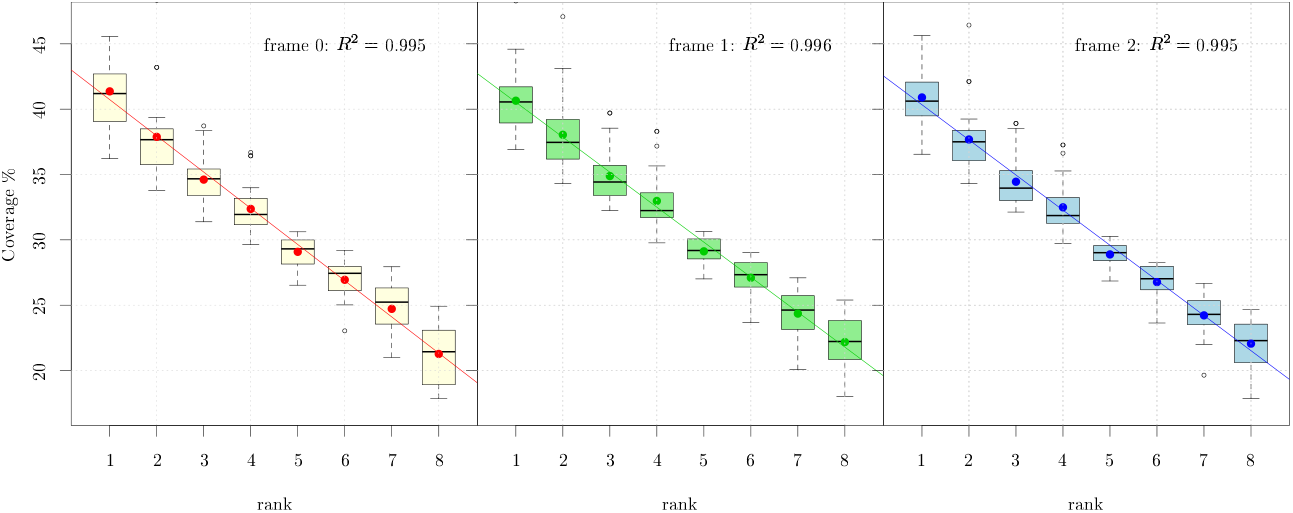
Universal scaling properties of the coverage within equivalence classes for the three reading frames.

In conclusion, each circular code has a distinct degree of coverage with respect to the species-specific codon usage of different organisms. Notably, however, behind this variability we observed universal properties, linking the coverage inside equivalence classes with the set of chemical transformations of the codons of the codes. Such strong organization is present in coding sequences but not in introns.

### 2.3. *Circular codes and* in vivo *translation speed*

The organization, present in the three frames of coding sequences and absent in intron sequences, hints at a biological role in the translation process. We explored this possibility by analysing the single codon translation speeds resulting from *E. coli his* operon attenuator reporter system Chevance et al. (2014). In this system, higher transcription rates of the reporter correspond to lower translation speeds. Remarkably, all the codons of the best code *X*_173_ fall within the set of fast translated codons, whereas the great part of the codons of code *X*_192_ appears to be among the slowest (see Figure 2). In order to verify whether this property holds for all the 27 equivalence classes we have computed the average speed for each code (i.e. the average speed of the set of 20 codons that compose each code) as a function of the coverage of the code in *E. coli* (i.e. the cumulative codon usage of the 20 codons of each code). Figure 3 shows the average speed versus the coverage for the 216 circular codes, where we have marked in blue the 27 codes that rank first within their equivalence class and in red the 27 codes that rank last. In order to enhance the comprehension we have reversed the scale so that higher values correspond to higher speeds. The best and worst codes form clusters that contain the fastest and the slowest codes, respectively. As mentioned above, the two sets are related by the chemical KM transformation. The relationship between circular-code-coverage and speed of translation appears to be linear with a correlation coefficient of 0.835. This would indicate that the coverage of a circular code can be a predictor of the speed of translation.

**Figure 2:**
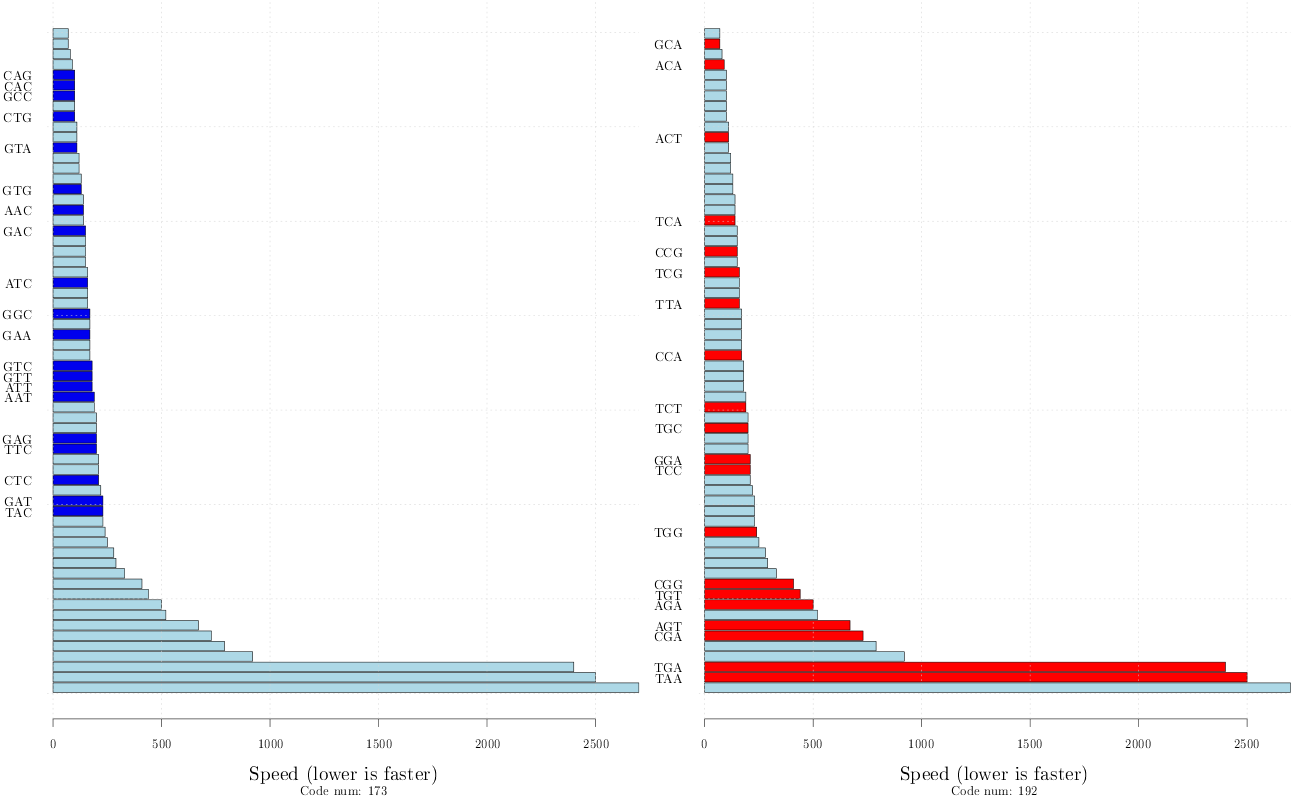
Ordered speed of the 64 codons, the data come from the experiment of Chevance et al. (2014) and lower values indicate faster codons. The codons coloured in blue(left) and in red(right) belong to code *X*_173_ and *X*_192_, respectively. They are the best and worst codes within the set of 8 codes forming the equivalence class shown in Table 1.

**Figure 3:**
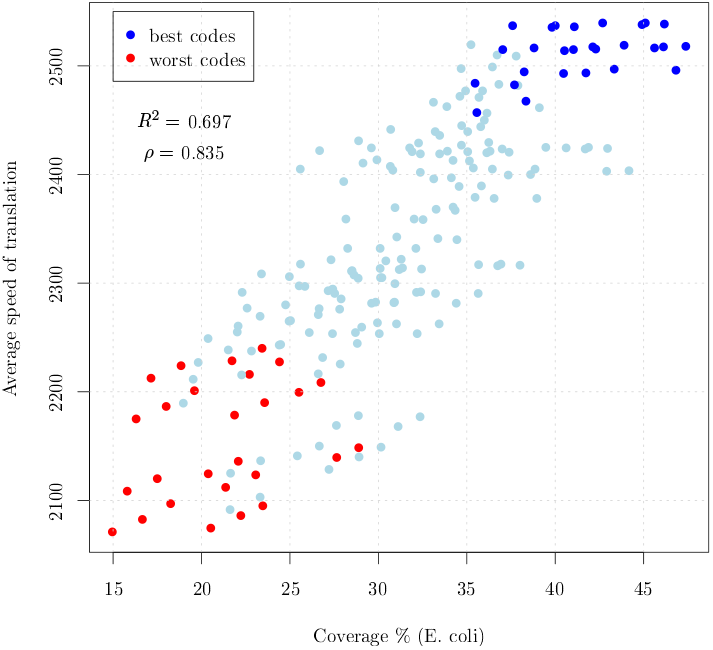
Average speed of translation versus Coverage (percent) for the 216 circular codes partitioned in 27 equivalence classes of 8 codes each. The points in blue and red correspond to the 27 best and 27 worst codes within their associated equivalence class. Clearly, the coverage is a predictor of the speed of translation and the best and worst codes within their equivalence class clusterize.

### 2.4. Circular codes and codon influence on protein expression

In order to further establish a link between circular codes theory and protein expression we analyzed the experimental evidence reported in Boël et al. (2016) where the authors use a black box logistic regression model over a large-scale protein expression dataset. Their aim was to assess the influence on protein expression of both mRNA sequence parameters and single codons. After accounting for sequence parameters such as predicted free folding energy or head folding indicators, they found a significant effect of individual codons that appears several positions after the initiator codon and stabilizes after about 16 codons. Conveniently, this analysis does not suffer from the presence of stop codons in the codes that may bias the average translation speed presented in Figure 3.

Consistently with the codon speed reported in the previous section, the codon influence is strongly correlated with the circular code coverage (*ρ* = 0.847, Figure 4). Notice that this cannot be explained in terms of single codon usage. Indeed, there is no evident correlation between single codon influence and single codon usage (Figure S2).

**Figure 4:**
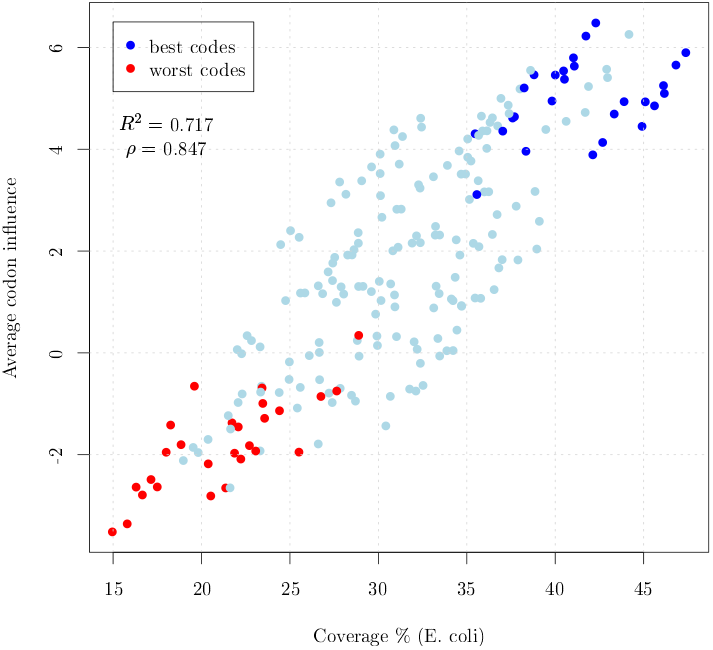
Average codon influence versus Coverage (percent) computed on the 216 circular codes partitioned in 27 equivalence classes of 8 codes each. The points in blue and red correspond to the 27 best and 27 worst codes within their associated equivalence class, respectively. As for the speed of translation (Figure 3), the coverage is a predictor of codon influence and the best and worst codes within their equivalence class clusterize.

### 2.5. Circular code motifs are absent in the mRNA 5’-head and 3’-tail sequences

Several independent reports demonstrated that the folding energy at the 5’ end of the mRNA explains most of the variation in protein expression levels, indicating that tightly folded messengers, obstructing the 30 nt ribosome binding site centered on the initiator codon, strongly influence translation initiation rates Kudla et al. (2009); Goodman et al. (2013); Cambray et al. (2018). In Boël et al. (2016) it is shown that, by computing the increase in the likelihood ratio when adding to the model terms corresponding to the average value of the codon influence over rolling windows of 5, 10 and 15 codons, the influence of codons is enhanced in the first part of the sequence, especially from codon 7 to codon 16 and stabilizes after 32-35 codons. If circular code properties play a role in translation, then we could expect a different coverage as a function of the position along the coding sequence. In Figure 5(left) we plotted the coverage of codes *X*_173_ (blue solid line) and *X*_192_ (red solid line) over rolling windows of 5 codons, computed over the first 100 codons of each complete coding sequence of *E.coli.* Remarkably, both for code *X*_173_ and *X*_192_ there is a transient initial span (around 40 codon positions) after which the rolling coverage over 5 codons reaches the value of the global coverage over the entire genome and fluctuates around it. While for code *X*_173_ the rolling coverage for the first positions is always lower than the global coverage, the rolling coverage for code *X*_192_ starts at a higher level with respect to the global coverage and decreases towards it. This appears to be a universal feature shared by all the organisms (see the Supplementary Information). The same is true for rolling windows up to 30 codons with no significant differences. The effect of the total codon content in the 3’ tail of the mRNA sequence was also reported to be influential on expression (Boël et al., 2016). Accordingly, we also observed a tail effect in the coverage of coding sequences (Figure 5(right) and Supplementary Information).

**Figure 5:**
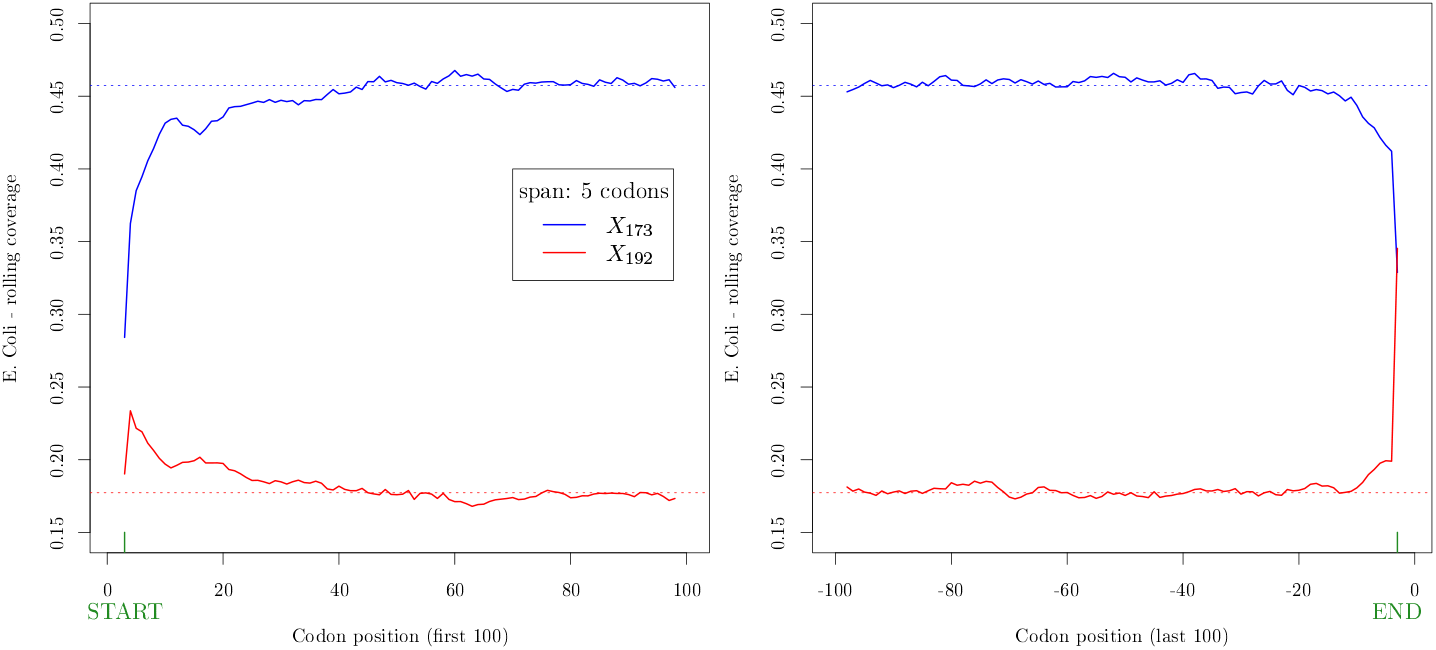
Rolling coverage (span: 5 codons) computed on the first (left) and last (right) 100 codon positions, averaged over the whole set of 3983 complete coding sequences of *E.coli*. The blue and red solid lines correspond to code *X*_173_ and *X*_192_, respectively. The dotted lines correspond to the global coverage of the codes over the whole genome.

These results indicate a lower coverage of the best circular code both in the head and in the tail of coding sequences, consistent with growing experimental evidence that other factors, such as mRNA folding energy, may predominate in those regions.

### 2.6. Protein expression levels correlate with circular code properties

To further explore the existence of a link between gene expression levels and circular codes, we computed the influence of each codon in a given sequence as the usage of such codons weighted by their specific influence. We analysed the set of 6348 sequences for which the protein expression levels had been previously measured and categorised, from 1 (low) to 5 (high) (Boël et al., 2016). This enabled us to correlate the expression level with the average cumulative influence of codons belonging to the best and worst codes (Figure 6). Clearly, a strong positive correlation (*ρ* = 0.93) between expression levels and the influence of the best code emerges (*ρ* = −0.97). Moreover, a strong negative correlation links the influence of the worst code to protein expression levels. Even more remarkably, the remaining codons (the 21 codons that do not belong to either of the two former codes) fail to show any noticeable correlation, so that, on average, an increase in the expression level score is linked to an increase of the circular code influence for the best code and a corresponding decrease for the worst code. In this way, a clear link between circular code properties and protein expression levels has been established, pointing to the existence of a role played by circular code properties in translation. As such, we anticipate that circular code theory can be important for the optimization of gene sequences for the production of recombinant proteins.

**Figure 6:**
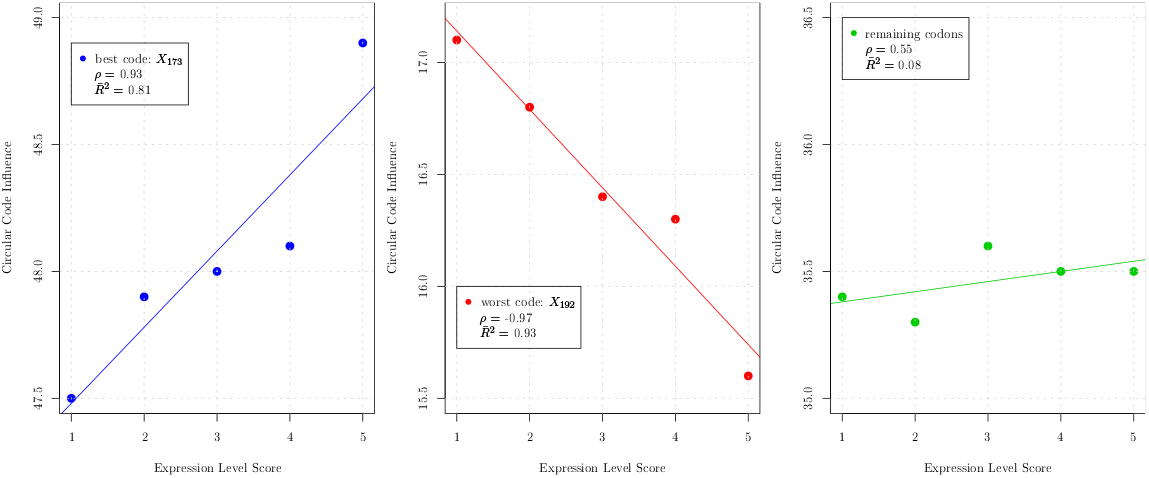
Expression level score vs average cumulative influence of circular codes. Left panel: best code *X*_173_. Center panel: worst code *X*_192_. Right panel: remaining codons, excluding stops.

### 2.7. circular code properties correlate with the S/W character of the first two nucleotides of the codon

Within each equivalence class the KM transformation always corresponds to passing from the best to the worst code, both in terms of coverage and translation efficiency, in agreement with recent experimental evidence of a correlation between a high codon usage and a high rate of decoding Gardin et al. (2014). In the KM transformation, keto (K; T or G) is transformed into amino (M; C or A) and viceversa (T↔C, G↔A). Hence, the KM transformation invariably changes the character of the base from strong (S; G or C) to weak (W; A or T), and this transformation appears to accompany remarkable effects on translation. Our results therefore indicate that the molecular biology in the decoding process may be significantly affected by the S/W character of the codon bases. Indeed, it has been reported that AT-rich codons are decoded slightly quicklier than GC-rich codons Gardin et al. (2014); Boël et al. (2016). AT-rich codons result in weaker secondary structures in mRNAs and therefore in higher translation initiation rates Goodman et al. (2013). However, at the elongation level a mechanistic explanation for faster decoding of AT-rich codons is still missing to date.

From a molecular point of view, an exact Watson-Crick base-pairing between codon and anticodon in the first two codon positions is indispensable for the correct decoding in the A-site of the ribosome Schmeing & Ramakrishnan (2009); Demeshkina et al. (2012). Moreover, functional and structural evidences indicate that during the decoding process universally conserved bases of the 16S rRNA closely interact with the codon-anticodon base-pair geometry in these positions Ogle et al. (2001). In particular, A1492 and A1493 adenosines form locally a triplex structure with the minor-groove of the codon-anticodon mini-helix (A-minor motif). These interactions appear to control domain closure of the 30S subunit Ogle et al. (2002), accelerating the forward steps in decoding, thus influencing the dynamics of translation elongation (recently reviewed in Opron & Burton (2019)). The evidence of minor-groove readout of the codon-anticodon mini-helix by the 16S rRNA A1492-A1493 dinucleotide bears interesting implications: because of nucleoside biochemistry, weak (W) basepairs (either A-U or U-A) have the same electron acceptor/donor profile in the minor groove. A-U or U-A are indistinguishable one from another with respect to the formation of an A-minor motif. The same applies for strong (S) basepairs: C-G or G-C display a different profile of electron donor/acceptors with respect to weak base pairs, but are indistinguishable one from another in the minor groove Masliah et al. (2013). Thus, out of the four different possible base pairs of two RNA nucleosides, only two possible hydrogen-bonding signatures can be discriminated in the minor groove, either weak (W) or strong (S).

In this respect, a striking feature, emerging from the analysis of the best and worst codes (e.g. *X*_173_ and *X*_192_, respectively), concerns the chemical nature of the bases of the first two nucleotides in the codon (Table 3). All the most influential codons of code *X*_173_ are of the kind SWN (strong-weak-any), the remaining ones being of the kind WWN (weak-weak-any). Conversely, by virtue of the KM transformation linking the two codes, the codons of code *X*_192_ are of the kind WSN or SSN. On average, these codons appear to be less influential. Hence, we investigated whether this property holds also for the remaining codes. We computed the average frequencies of SWN, WWN, SSN and WSN codons for the group of best codes (blue) and worst codes (red), see Figure S5, where the area of the bubbles is proportional to the average influence of each group of codons. Clearly, codons of the kind SWN and WWN identify the best codes, i.e. those associated to a higher expression level and coverage. Conversely, codons of the kind SSN and WSN characterize the codes having lower expression level and coverage.

**Table 3:**
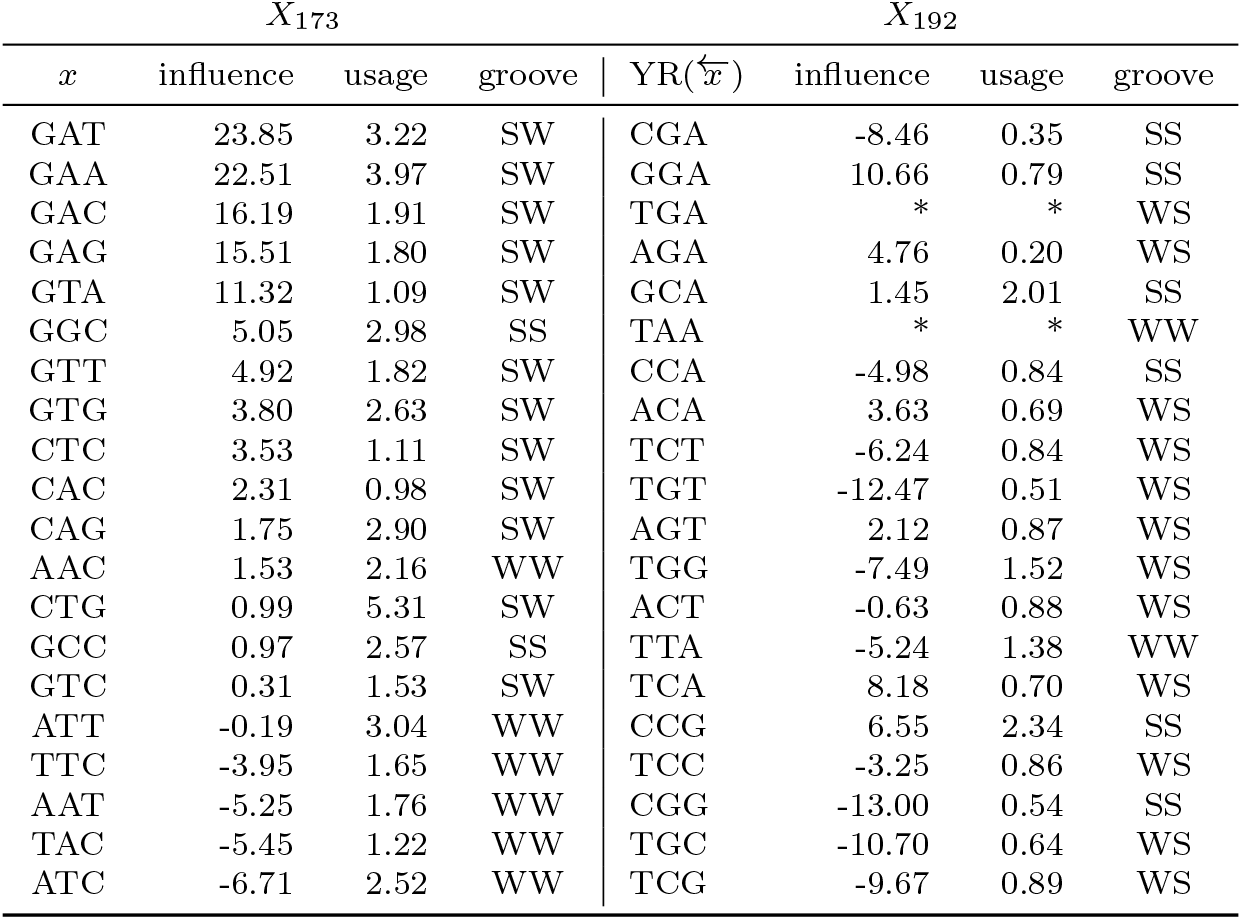
Codons of circular codes X_173_ and X_192_ together with their codon influence as in Boёl et al. (2016) their codon usage in E.coli and the mRNA groove described as the Strong/Weak nature of the first two nucleotides of the codon. The columns are ordered in descending order according to the codon influence index for code X_173_ (second column).

Hence, if the A-minor motif forms a structure able to monitor the correct base-pairing of the first two bases of the codon, through readout of the minor groove, then the dichotomic combination of S/W base pairs in these positions may impose different spatial arrangements of the 16S rRNA through A1492-A1493 dinucleotide interaction influencing the speed/rates of mRNA decoding by the ribosome. Strikingly, the analysis of circular code properties appears to point at a link between the S/W dichotomy in the first two bases of the codon and protein expression levels. In particular, the results indicate a codon ordering where SWN codons confer the highest expression levels. In this respect, the theory of circular codes allowed to uncover the possible role played by the S/W dichotomy in the decoding process. It is also worth noticing that without the lens of circular codes this property would have otherwise escaped from the analysis of synonymous sequence libraries, since the latter tend to vary mostly in the third (wobbling) position of the codon, and only marginally in the first two positions (only for degeneracy-6 codons).

## 3. Conclusions

We have shown that circular codes theory provides a new and powerful key to understanding the influence of codon bias on gene expression. Circular code coverage exhibits taxon-independent universal properties with a strong hierarchical organization. Independently from codon usage, universal frame marks are present in coding sequences and are absent in introns. Indeed, there are recurring properties, linking the coverage inside equivalence classes with the set of chemical transformations of the codons of the codes. These properties strongly correlate with translation speed, codon influence and protein expression level. In accordance with the predominant effect of the secondary structure of mRNAs in the 5’ ends on translation, circular code properties are absent at the beginning of coding sequences, and correlate with the S/W dichotomy in the first two nucleotides of codons.

For these reasons the theory of circular codes can be also seen as a promising tool for codon optimization of protein coding sequences to be used in biotechnological applications and for building sequence indicators for bioinformatics applications. If circular code properties play a role in translation then it will be possible to design dedicated experiments to verify their impact on expression rates and/or reading frame maintainance paving the way to a better understanding of the molecular mechanisms behind decoding.

## Author Contributions

All the authors contributed equally to this study.

