## Supplementary Information for "A role for circular code properties in translation"

### Supplemental Information for:
### A role for circular code properties in translation

Simone Giannerini[a,*], Alberto Danielli[b], Diego Luis Gonzalez[a,c], Greta Goracci[a]

[a] *Department of Statistical Sciences, University of Bologna, Bologna, 40126, Italy.*
[b] *Department of Pharmacy and Biotechnology, University of Bologna, Bologna, 40126, Italy.*
[c] *Institute for Microelectronics and Microsystems - Bologna Unit, CNR, Bologna, 40129, Italy.*

**Abstract**

This supplement follows the same sectioning of the article. In the first section we present a minimal introduction to comma-free codes and circular codes. In the second section we report the extended results for the analysis presented in the main article.

#### 1. Circular codes and comma free codes

In analogy with the transmission of a digital message, an efficient protein synthesis needs appropriate means to achieve the following fundamental tasks: *i*) determine the points where translation should start and stop, *ii*) avoid reading errors due to frame shifts, that is, ensure that the ribosome stays synchronized with the correct reading frame. The latter ability is called *reading frame maintenance* and is crucial since an error would result in a completely wrong protein. While the problem of punctuation signs has been elucidated to a great extent, the problem of explaining the determinants of frame maintenance are still largely unknown. As mentioned in the Introduction, reading frame synchronization in mRNA translation was first studied in Crick et al. (1957), which proposed an elegant solution based on comma free codes. A **comma free** code is a special set of codons that allows to retrieve the normal reading frame anywhere in the sequence, provided this is composed only of codons of such code. The idea can be explained by means of the following simple example:

**Example 1.** The comma free code $X$ has two codons

$$X = \{\text{CTG}, \text{AAT}\}$$

1. Build a sequence with the codons of $X$ (in green), for instance

$$\text{AAT CTG AAT AAT}$$

2. Read it in the 3 possible frames:

frame 0: AAT CTG AAT AAT

frame 1: ~~A~~ ATC TGA ATA ~~AT~~

frame 2: ~~AA~~ TCT GAA TAA ~~T~~

3. There is only one frame (frame 0) where all the codons belong to $X$: the *correct reading frame*. **None** of the codons (in red) read in frames +1 and +2 belong to $X$.

This holds for any sequence of arbitrary length formed with codons of $X$.

In other words, if we form sequences by using codons of a comma free code and we read them with a frame shift then we end up immediately on a *forbidden* codon, i.e. a codon that does not belong to the code. Despite their appeal, comma free codes were proven not adequate and left aside, especially after the experiment of Nirenberg & Matthaei (1961), which showed that the codon TTT codes for the amino acid Phenylalanine but, for theoretical reasons, TTT cannot be a part of a any comma-free code. In general, one can argue that it is not possible to identify good codons and build a code with them since all the 64 codons are used in protein synthesis; there are no forbidden or bad codons.

After 40 years from Crick's proposal of comma-free codes, Arquès & Michel (1996) found empirically that a weaker version of comma-free codes can be used to retrieve the normal reading frame. These are called *circular codes* and can be explained through the following example:

**Example 2.** Assume that the circular code $X$ has 3 codons

$$X = \{\text{CTG}, \text{AAT}, \text{TGA}\}.$$

1. Form an arbitrary sequence with the codons of $X$, for instance:

CTG AAT CTG

2. Put it in a circle and read it in the 3 possible frames (the starting nucleotide is coloured in blue):

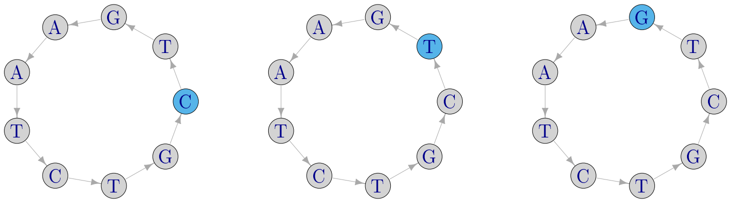

frame 0 : CTG AAT CTG    frame 1 : TGA ATC TGC    frame 2 : GAA TCT GCT

3. There is only one frame (frame 0) where all the codons belong to $X$: the *correct reading frame*, even if some of the codons read in frames +1 and +2 can belong to $X$.

| | | | |
|---|---|---|---|
| 1. (A)(T)(C)(G) | : A ↦ A; T ↦ T; C ↦ C; G ↦ G | Identity | (I) |
| 2. (AT)(CG) | : A ↦ T; T ↦ A; C ↦ G; G ↦ C | Strong/Weak | (SW) |
| 3. (AG)(CT) | : A ↦ G; G ↦ A; C ↦ T; T ↦ C | Purine/Pyrimidine | (YR) |
| 4. (AC)(GT) | : A ↦ C; C ↦ A; G ↦ T; T ↦ G | Keto/Amino | (KM) |
| 5. (A)(T)(CG) | : A ↦ A; T ↦ T; C ↦ G; G ↦ C | | |
| 6. (AT)(C)(G) | : A ↦ T; T ↦ A; C ↦ C; G ↦ G | | |
| 7. (ACTG) | : A ↦ C; C ↦ T; T ↦ G; G ↦ A | | |
| 8. (AGTC) | : A ↦ G; G ↦ T; T ↦ C; C ↦ A | | |

Table 1: Set of the 8 transformations of the nucleotides forming the dihedral group $D_8$. The first 4 transformations of the nucleotides form also the Klein V symmetry group. These are indicated as chemical transformations in the rightmost column.

As before, this holds for any sequence of arbitrary length formed with codons of $X$. The two examples pinpoint the difference between comma-free and circular codes: for comma-free codes a frame shift in the sequence invariably leads to a forbidden codon, whereas for circular codes this is not necessarily so and valid codons can be found when the sequence is read out of frame.

It is easy to show that a circular code can have at most 20 codons. If this is the case, the code is said *maximal*. The codes found in Arquès & Michel (1996) are maximal and have two additional properties: 1) they are *self complementary*: if a codon belongs to a code, then also its reverse complement belongs to the code; 2) they are $C^3$: the circular permutations of the codons of a circular code also form a circular code. We denote with $\alpha_1(x)$ and $\alpha_2(x)$ the two circular permutations of a codon $x$. For example, if $x = \text{CTG}$ then $\alpha_1(x) = \text{TGC}$ and $\alpha_2(x) = \text{GCT}$. There are exactly 216 circular codes that possess the three aforementioned properties i.e. they are maximal, self-complementary and $C^3$. In the following we use the list of 216 codes given in Michel et al. (2008) and label them according to the order given there, so that we denote a generic $i$-th code of 20 codons with $X_i$, $i = 1, \ldots, 216$. In Fimmel et al. (2015) we have shown that these 216 codes have special symmetry properties related to the transformations of the nucleotides. A transformation is a rule that maps the set of 4 nucleotides onto one of its 24 possible permutations. For instance, the transformation (AGT)(C) maps A to G, G to T, T to A and C to C, that is

$$\text{A} \mapsto \text{G}; \quad \text{G} \mapsto \text{T}; \quad \text{T} \mapsto \text{A}; \quad \text{C} \mapsto \text{C}\,.$$

There are 8 special transformations that are related to the dihedral group of symmetry, that is, they represent the 8 symmetries of a square. These are shown in Table S1. Note that a single letter within brackets means that it is not transformed and for ease of notation it can be omitted. Hence, in the example above, (AGT)(C) becomes (AGT). The first four transformations of the list form a further group of symmetry (the Klein V group) that contains the identity plus three chemical transformations of the nucleotides (Gonzalez et al., 2008). An important result proved in Fimmel et al. (2015) states that by means of the 8 above transformations it is possible to partition the 216 codes in

| | I $X_{173}$ | (AT) $X_{176}$ | (CG) $X_{203}$ | SW $X_{206}$ | YR $X_{183}$ | (ACTG) $X_{182}$ | (AGTC) $X_{193}$ | KM $X_{192}$ |
|---|---|---|---|---|---|---|---|---|
| 1 | AAC | TTC | AAG | TTG | GGT | GGA | CCT | CCA |
| 2 | GTT | GAA | CTT | CAA | ACC | TCC | AGG | TGG |
| 3 | AAT | TTA | AAT | TTA | GGC | GGC | CCG | CCG |
| 4 | ATT | TAA | ATT | TAA | GCC | GCC | CGG | CGG |
| 5 | ATC | TAC | ATG | TAG | GCT | GCA | CGT | CGA |
| 6 | GAT | GTA | CAT | CTA | AGC | TGC | ACG | TCG |
| 7 | CAC | CTC | GAG | GTG | TGT | AGA | TCT | ACA |
| 8 | GTG | GAG | CTC | CAC | ACA | TCT | AGA | TGT |
| 9 | CAG | CTG | GAC | GTC | TGA | AGT | TCA | ACT |
| 10 | CTG | CAG | GTC | GAC | TCA | ACT | TGA | AGT |
| 11 | CTC | CAC | GTG | GAG | TCT | ACA | TGT | AGA |
| 12 | GAG | GTG | CAC | CTC | AGA | TGT | ACA | TCT |
| 13 | GAA | GTT | CAA | CTT | AGG | TGG | ACC | TCC |
| 14 | TTC | AAC | TTG | AAG | CCT | CCA | GGT | GGA |
| 15 | GAC | GTC | CAG | CTG | AGT | TGA | ACT | TCA |
| 16 | GTC | GAC | CTG | CAG | ACT | TCA | AGT | TGA |
| 17 | GCC | GCC | CGG | CGG | ATT | TAA | ATT | TAA |
| 18 | GGC | GGC | CCG | CCG | AAT | TTA | AAT | TTA |
| 19 | GTA | GAT | CTA | CAT | ACG | TCG | AGC | TGC |
| 20 | TAC | ATC | TAG | ATG | CGT | CGA | GCT | GCA |

Table 2: Equivalence class formed by eight circular codes. Each column represents one of the 216 circular codes. The codes are related through the group of transformations $D_8$. For instance AAC $\in X_{173}$ and KM(AAC) = CCA $\in X_{192}$.

27 equivalence classes, see Table S2. Each equivalence class contains 8 circular codes related through the 8 transformations of the dihedral group shown in Table S1. Formally, two circular codes $X_j$ and $X_z$ are equivalent iff there exists a transformation $\pi$ of the group $D_8$ such that $X_z = \pi(X_j)$. The classification in equivalence classes is one of the key aspects connecting the theory of circular codes with the experimental results on protein expression. In Table S2 we show an example of one of the 27 equivalence classes. For instance, from the first row of the table we can see that codon AAC belongs to code number 173 $X_{173}$) whereas its Keto-Amino transformation KM(AAC) = CCA belongs to code 192 ($X_{192}$) and so on.

##### *1.1. Coverage of a circular code*

The **coverage** of a circular code over a specific sequence or organism, is a key quantity to study the role played by circular codes in translation. It is the cumulative codon usage of the codons belonging to a code and can be seen as a measure of the "goodness" of a code, see also Gonzalez et al. (2011). It can be interpreted as a sort of aggregate codon usage of the set of codons of the code. In the following we provide a rigorous mathematical definition.

Given a genome $i$, we define its codon distribution (or codon usage) $\mathbf{p}_i$ over the set of codons of $\mathcal{B}^3$ as:

| Codons | $x_{1i}$ | $\ldots$ | $x_{ki}$ | $\ldots$ | $x_{64i}$ |
|---|---|---|---|---|---|
| Usage | $p_{1i}$ | $\ldots$ | $p_{ki}$ | $\ldots$ | $p_{64\,i}$ |

| | I | (AT) | (CG) | SW | YR | (ACTG) | (AGTC) | KM |
|---|---|---|---|---|---|---|---|---|
| **1** | 173 | 176 | 203 | 206 | 183 | 182 | 193 | 192 |
| **2** | 23 | 33 | 77 | 81 | 13 | 37 | 65 | 87 |
| **3** | 98 | 10 | 96 | 8 | 52 | 45 | 55 | 53 |
| 4 | 25 | 35 | 76 | 85 | 50 | 47 | 59 | 56 |
| 5 | 20 | 34 | 75 | 80 | 17 | 40 | 69 | 89 |
| **6** | 166 | 216 | 164 | 213 | 186 | 187 | 189 | 191 |
| **7** | 4 | 104 | 6 | 102 | 16 | 42 | 61 | 86 |
| 8 | 27 | 30 | 72 | 84 | 38 | 12 | 88 | 64 |
| 9 | 117 | 160 | 118 | 157 | 130 | 131 | 133 | 135 |
| 10 | 111 | 159 | 116 | 151 | 119 | 126 | 138 | 145 |
| 11 | 22 | 29 | 71 | 79 | 2 | 1 | 100 | 99 |
| **12** | 172 | 175 | 202 | 205 | 181 | 184 | 196 | 195 |
| **13** | 21 | 31 | 74 | 78 | 11 | 39 | 68 | 91 |
| **14** | 24 | 32 | 73 | 83 | 49 | 48 | 60 | 57 |
| **15** | 97 | 9 | 95 | 7 | 51 | 46 | 58 | 54 |
| **16** | 171 | 174 | 201 | 204 | 167 | 178 | 200 | 208 |
| **17** | 3 | 103 | 5 | 101 | 15 | 43 | 62 | 90 |
| **18** | 165 | 215 | 163 | 212 | 185 | 188 | 190 | 194 |
| 19 | 26 | 28 | 70 | 82 | 36 | 14 | 92 | 66 |
| 20 | 123 | 124 | 141 | 143 | 105 | 106 | 150 | 147 |
| **21** | 115 | 158 | 113 | 155 | 129 | 132 | 134 | 136 |
| **22** | 161 | 214 | 162 | 211 | 168 | 179 | 197 | 207 |
| 23 | 122 | 125 | 140 | 142 | 110 | 108 | 152 | 149 |
| **24** | 41 | 94 | 18 | 67 | 19 | 44 | 63 | 93 |
| **25** | 107 | 156 | 112 | 148 | 120 | 127 | 139 | 146 |
| 26 | 177 | 210 | 169 | 199 | 170 | 180 | 198 | 209 |
| 27 | 137 | 121 | 144 | 128 | 114 | 109 | 153 | 154 |

Table 3: List of the 216 maximal, self-complementary, $C^3$ circular codes partitioned into 27 equivalence classes. Each class contains 8 codes linked through the chemical transformations of the dihedral group $D_8$. We highlighted in bold the 16 classes for which the codes corresponding to the identity (I, first column) and to the Keto-Amino transformation (KM, last column) have no common codons (they are disjoint).

where $x_k \in \mathcal{B}^3$ and $p_{ki} \in \mathbf{p}_i$. Next, we define the coverage of a code as the cumulative codon usage over the set of codons that compose the code.

**Definition 3.** Given a circular code $X_j \in \mathfrak{C}$ where $j = 1, \ldots, 216$ and a genome $i$, we define as $\mathrm{C}_{ij}$ the coverage of code $X_j$ over genome $i$:

$$\mathrm{C}_{ij} = \sum_{k=1}^{64} p_{ki} \, I_{X_j}(x_k) \tag{1}$$

where $I_A(x)$ is the indicator function, i.e.

$$I_A(x) = \begin{cases} 1, & \text{if } x \in A \\ 0, & \text{if } x \notin A. \end{cases}$$

Clearly, the coverage ranges in $[0, 1]$.

**Example 4.** Consider the sequence CAT CTG AAT GGA CTG and the two codes $X_1 = \{\mathrm{CTG}, \mathrm{AAT}\}$, $X_2 = \{\mathrm{GGA}, \mathrm{TGT}\}$. The codon usage of the sequence is

| Codons | CAT | CTG | AAT | GGA |
|---|---|---|---|---|
| Usage | 1/5 | 2/5 | 1/5 | 1/5 |

The coverage of $X_1$ results $2/5 + 1/5 = 3/5 = 0.60$, and that of $X_2$ results $1/5 = 0.20$.

#### 2. Results

##### *2.1. Universal properties of Circular codes' coverage*

We have analyzed the whole Codon Usage Database available at `http://www.kazusa.or.jp/codon/` Nakamura et al. (1997). It contains 35799 organisms and 3,027,973 complete protein coding genes (CDS). After some cleaning and removing the mitochondrial genomes we end up with 25528 nuclear genomes. In Table S4 we report a brief summary of the database. In Table 2 of the main article we show the coverage (in percentage), the rank over the 216 codes and the rank within a class for the equivalence class composed of the 8 codes shown in Table S2. The ranks inside the equivalence class show a universal ordering among the 8 codes, irrespective of the species-specific codon usage. In particular, the worst code within each class (code with the least coverage) invariably coincides with the chemical Keto-Amino transformation of the best one. As we will show, this property holds for all the equivalence classes. Here and in the following we focus on the 16 equivalence classes for which the best and the worst codes (w.r.t coverage) are disjoint sets.

| species | # organisms |
|---|---|
| bacteria | 4918 |
| animals | 6921 |
| viral | 1956 |
| plants | 11733 |
| total | 25528 |

| domain | # organisms |
|---|---|
| prokaryotes | 4918 |
| eukaryotes | 15598 |
| total | 20516 |

Table 4: Description of the codon usage database analyzed. The right table shows the aggregation by domain, excluding viruses and phage.

The results shown in Table S5 demonstrate that universal symmetry properties of coding sequences emerge when analyzed through the theoretical framework of circular codes, irrespectively of the species-specific codon-usage. Moreover, within each equivalence class, the Keto-Amino transformation of the code possessing the best coverage always leads to the worst covering code of the same class. This establishes an important connection between the codon usage, the Keto-Amino (KM) and the Purine-Pyrimidine (YR) chemical transformation. Behind the heterogeneity of the codon usage, there is a universal ordering structure conserved across domains of life, grounded by a theoretical framework of circular codes. The aforementioned properties emerge only if we consider these special set of codons (codes) and does not necessarily hold at the level of single codons. This is substantiated further by means of a statistical test.

*2.1.1. A bootstrap test*

In this section we present a bootstrap test to explore the relation between the coverages of the best and the worst code of an equivalence class. In the previous section we have shown that, within each equivalence class, the code $X \in \mathfrak{C}$ that has the best coverage is (almost) unique. Also, the code that has the worst coverage is $\mathrm{KM}(X)$, the Keto-Amino (Rumer) transformation of code $X$. Is this result due to sheer chance?

It is natural to expect that the more a set of codons is recurrent then the less recurrent are codons that do not belong to that set. We have selected from the database the genomes with at least 1 million codons. There are 291 such genomes on which we have computed the coverage of the pairs of codes $X_{173}, X_{192}$ and $X_{23}, X_{87}$. These are shown in Figure 1, where we have superimposed the following quadratic least square fits (blue points):

$$C_{i,192} = \beta_{0,173} + \beta_{1,173} C_{i,173} + \beta_{2,173} C_{i,173}^2 + \varepsilon_i$$
$$C_{i,87} = \beta_{0,23} + \beta_{1,23} C_{i,23} + \beta_{2,23} C_{i,23}^2 + \varepsilon_i$$

where $i = 1, \ldots, 291$ genomes. Clearly, the quadratic fit accounts for 87% of the observed variability. Hence, the question is: is this negative correlation due to the fact that the more one set is recurrent, then the less recurrent is its complement? In other words, is this correlation compatible with the natural correlation produced by a random choice of codons? For instance, in the equivalence class coverage

| best | $X_{173}$ | $X_{23}$ | $X_{98}$ | $X_{166}$ | $X_{4}$ | $X_{172}$ | $X_{21}$ | $X_{24}$ | $X_{97}$ | $X_{171}$ | $X_{3}$ | $X_{165}$ | $X_{115}$ | $X_{161}$ | $X_{41}$ | $X_{107}$ |
|---|---|---|---|---|---|---|---|---|---|---|---|---|---|---|---|---|
| bacteria | 46.4 | 46.5 | 46.0 | 44.3 | 44.4 | 43.5 | 43.7 | 42.9 | 43.2 | 43.0 | 41.6 | 41.4 | 40.8 | 40.9 | 39.0 | 38.6 |
| animals | 42.0 | 41.3 | 41.6 | 41.3 | 40.6 | 40.0 | 39.3 | 40.1 | 39.6 | 39.4 | 38.6 | 39.3 | 39.4 | 38.7 | 37.9 | 36.4 |
| viral | 43.2 | 42.7 | 42.0 | 42.6 | 42.1 | 41.0 | 40.5 | 41.1 | 39.8 | 40.6 | 39.9 | 40.4 | 40.5 | 40.0 | 38.7 | 37.5 |
| plants | 39.7 | 40.1 | 39.6 | 39.6 | 40.0 | 40.0 | 40.5 | 40.4 | 39.9 | 40.8 | 40.3 | 39.9 | 40.3 | 40.7 | 38.8 | 38.6 |
| worst | $X_{192}$ | $X_{87}$ | $X_{53}$ | $X_{191}$ | $X_{86}$ | $X_{195}$ | $X_{91}$ | $X_{57}$ | $X_{54}$ | $X_{208}$ | $X_{90}$ | $X_{194}$ | $X_{136}$ | $X_{207}$ | $X_{93}$ | $X_{146}$ |
| bacteria | 18.1 | 19.7 | 19.9 | 17.1 | 18.7 | 17.0 | 18.6 | 20.5 | 18.8 | 22.1 | 17.6 | 16.0 | 19.5 | 21.1 | 22.7 | 23.5 |
| animals | 23.4 | 23.8 | 23.4 | 22.4 | 22.8 | 21.7 | 22.1 | 24.2 | 21.7 | 24.6 | 21.1 | 20.7 | 23.2 | 23.6 | 24.3 | 25.7 |
| viral | 22.4 | 22.5 | 22.8 | 21.6 | 21.7 | 20.4 | 20.5 | 23.5 | 20.9 | 23.6 | 19.7 | 19.6 | 22.7 | 22.8 | 23.7 | 25.0 |
| plants | 23.5 | 25.2 | 23.9 | 22.9 | 24.7 | 21.7 | 23.5 | 22.4 | 22.2 | 24.2 | 22.9 | 21.2 | 21.9 | 23.6 | 23.7 | 25.7 |

absolute rank

| best | $X_{173}$ | $X_{23}$ | $X_{98}$ | $X_{166}$ | $X_{4}$ | $X_{172}$ | $X_{21}$ | $X_{24}$ | $X_{97}$ | $X_{171}$ | $X_{3}$ | $X_{165}$ | $X_{115}$ | $X_{161}$ | $X_{41}$ | $X_{107}$ |
|---|---|---|---|---|---|---|---|---|---|---|---|---|---|---|---|---|
| bacteria | 2 | 1 | 3 | 7 | 6 | 16 | 13 | 19 | 17 | 18 | 22 | 24 | 29 | 28 | 35 | 39 |
| animals | 2 | 7 | 3 | 6 | 9 | 12 | 18 | 11 | 14 | 16 | 21 | 17 | 15 | 20 | 27 | 37 |
| viral | 2 | 4 | 9 | 6 | 8 | 12 | 15 | 11 | 22 | 14 | 20 | 17 | 16 | 19 | 29 | 35 |
| plants | 16 | 9 | 18 | 17 | 12 | 11 | 4 | 5 | 15 | 1 | 7 | 14 | 8 | 2 | 20 | 22 |
| worst | $X_{192}$ | $X_{87}$ | $X_{53}$ | $X_{191}$ | $X_{86}$ | $X_{195}$ | $X_{91}$ | $X_{57}$ | $X_{54}$ | $X_{208}$ | $X_{90}$ | $X_{194}$ | $X_{136}$ | $X_{207}$ | $X_{93}$ | $X_{146}$ |
| bacteria | 212 | 207 | 206 | 214 | 210 | 215 | 211 | 205 | 209 | 196 | 213 | 216 | 208 | 201 | 191 | 183 |
| animals | 208 | 205 | 207 | 211 | 210 | 214 | 212 | 202 | 213 | 200 | 215 | 216 | 209 | 206 | 201 | 189 |
| viral | 209 | 208 | 203 | 211 | 210 | 214 | 213 | 202 | 212 | 201 | 215 | 216 | 207 | 205 | 200 | 185 |
| plants | 208 | 195 | 202 | 210 | 198 | 215 | 209 | 212 | 213 | 201 | 211 | 216 | 214 | 205 | 203 | 187 |

relative rank

| best | $X_{173}$ | $X_{23}$ | $X_{98}$ | $X_{166}$ | $X_{4}$ | $X_{172}$ | $X_{21}$ | $X_{24}$ | $X_{97}$ | $X_{171}$ | $X_{3}$ | $X_{165}$ | $X_{115}$ | $X_{161}$ | $X_{41}$ | $X_{107}$ |
|---|---|---|---|---|---|---|---|---|---|---|---|---|---|---|---|---|
| bacteria | 1 | 1 | 1 | 1 | 1 | 1 | 1 | 1 | 1 | 1 | 1 | 1 | 1 | 1 | 1 | 1 |
| animals | 1 | 1 | 1 | 1 | 1 | 1 | 1 | 1 | 1 | 1 | 1 | 1 | 1 | 1 | 1 | 1 |
| viral | 1 | 1 | 1 | 1 | 1 | 1 | 1 | 1 | 1 | 1 | 1 | 1 | 1 | 1 | 1 | 1 |
| plants | 1 | 1 | 1 | 1 | 1 | 1 | 1 | 1 | 1 | 1 | 1 | 1 | 1 | 1 | 1 | 1 |
| worst | $X_{192}$ | $X_{87}$ | $X_{53}$ | $X_{191}$ | $X_{86}$ | $X_{195}$ | $X_{91}$ | $X_{57}$ | $X_{54}$ | $X_{208}$ | $X_{90}$ | $X_{194}$ | $X_{136}$ | $X_{207}$ | $X_{93}$ | $X_{146}$ |
| bacteria | 8 | 8 | 8 | 8 | 8 | 8 | 8 | 8 | 8 | 8 | 8 | 8 | 8 | 8 | 8 | 8 |
| animals | 8 | 8 | 8 | 8 | 8 | 8 | 8 | 8 | 8 | 8 | 8 | 8 | 8 | 8 | 8 | 8 |
| viral | 8 | 8 | 8 | 8 | 8 | 8 | 8 | 8 | 8 | 8 | 8 | 8 | 8 | 8 | 8 | 7 |
| plants | 8 | 8 | 8 | 8 | 8 | 8 | 8 | 8 | 8 | 8 | 8 | 8 | 8 | 8 | 8 | 8 |

Table 5: Coverage (upper panel), absolute ranks (mid panel) and relative ranks (lower panel) for the best and worst codes of the 16 equivalence classes highlighted in bold in Table S3 (the best and the worst codes are disjoint sets). The universality of the results is clear if we consider the ranks within classes: for instance the coverage of code $X_{173}$ ($X_{192}$) for bacteria is 46.4 (18.1) (upper panel). It is not the highest (lowest) among the 216 codes, indeed it ranks 2nd (212th) (mid panel). However, it is always the highest (lowest) within its class (lower panel).

.

of Table S2 the best and the worst codes are $X_{173}$ and $X_{192}$, respectively. Now, we show that $C_{i,192}$, i.e. the coverage of $X_{192}$ is significantly smaller than the coverage of a random set of 20 codons taken from those who do not belong to the best code $X_{173}$. Formally, denote with $\mathrm{C}_{ij}$ and $\mathrm{C}_{ij'}$ the coverage of the codes $X_j$ and $X_{j'} = \mathrm{KM}(X_j)$ over a genome $i$. Also, let $D = \mathcal{B}^3 \setminus X_j$ be the set of 44 codons that do not belong to $X_j$. In statistical terms the system of hypotheses results

$$\begin{cases} H_0 : \mathrm{C}_{ij} \text{ is compatible with } \mathrm{C}_D \\ H_1 : \mathrm{C}_{ij} \text{ is not compatible with } \mathrm{C}_D \end{cases} \tag{2}$$

where $\mathrm{C}_D$ is the following random variable: coverage of a random set of 20 codons taken from $D$. In order to test this hypothesis we implement the following bootstrap scheme:

1. Compute $\mathrm{C}_{ij}$ and $\mathrm{C}_{ij'}$ the coverage of the codes $X_j$ and $X_{j'} = \mathrm{KM}(X_j)$ over a genome $i$.
2. Resample without replacement B sets of 20 elements from $D = \mathcal{B}^3 \setminus X_j$ and compute the coverage over genome $i$ for each resample:

   | Set | $\bar{X}^*_{j1}$ | ... | $\bar{X}^*_{jB}$ |
   |---|---|---|---|
   | Coverage | $\bar{\mathrm{C}}^*_{ij1}$ | ... | $\bar{\mathrm{C}}^*_{ijB}$ |

   The set of resampled codes can be made homogeneous with respect to the GC content by imposing that their GC content be equal to that of the original code $X_{j'}$.
3. Compare the coverage $\mathrm{C}_{ij'}$ with the quantiles of the empirical distribution function of the coverage $\bar{F}^*_{ij}$. Alternatively, compute the bootstrap $p$-value $p = \frac{1}{B}\sum_{b=1}^{B} I(\bar{\mathrm{C}}^*_{ijb} > \mathrm{C}_{ijb})$.

At Step 2 of the algorithm the resampling of the sets of codons from $D$ can be performed in two ways: *i*) using a uniform distribution over $D$; *ii*) using the codon distribution of genome $i$ over $D$. The hypotheses can have a different biological meaning/interpretation. In brief, the first hypothesis assumes that all the 216 codes are equally likely to occur in practice and exist independently from the codon usage of the genomes. The second hypothesis, instead, assumes that the occurrence of a circular codes is related to the codon usage of a given genome. Since the results point to a universal relationship which is independent of the codon usage in genomes we tend to support the first hypothesis. In any case, we have performed the tests for both hypotheses and we show the results in Figure 1. The red lines correspond to the Monte Carlo rejection bands for hypothesis *i*): the occurrence of the codes is uniform; the green lines are the rejection bands for hypothesis *ii*): the occurrence of the codes depends on the codon usage. In both cases $\alpha = 0.001$ (i.e. $B = 9999$ bootstrap replicates). The results show clearly that in both scenarios $H_0$ is rejected so that the negative correlation observed cannot be ascribed to random fluctuations and goes well beyond the naturally induced correlation.

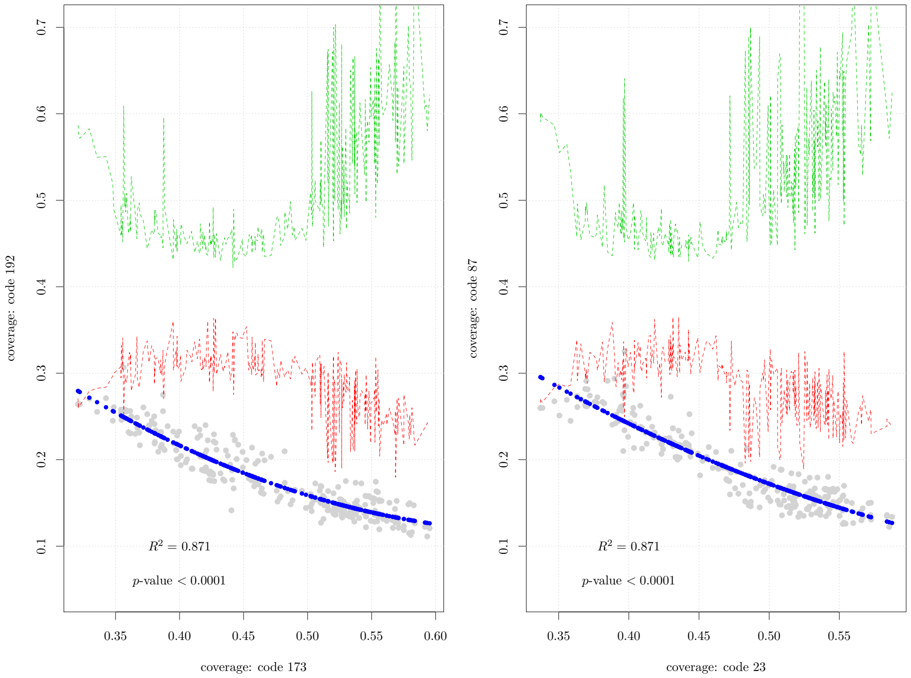

Figure 1: Coverage for the best and worst codes of two equivalence classes, computed over 291 genomes with more than 1 million codons (gray points). The blue points represent a quadratic fit and the associated $R^2$ and $p$-value are reported under it. The green and red lines correspond to the bootstrap rejection bands under the null hypothesis at $\alpha = 0.001$ that the observed relation is produced by chance, as follows: red line, hypothesis 1: the occurrence of the codes is uniform; green line, hypothesis 2: the occurrence of the codes follows the codon usage. In both cases the null hypothesis that the negative correlation is due to chance is clearly rejected.

*2.2. Universal frame marks in coding sequences.*

In this section we report the extended analysis in the three reading frames over the whole set of complete coding sequences for 25 organisms. The description of the genomes is reported in Table S6.

| organism | no. sequences | no. codons |
|---|---:|---:|
| AeropyrumPernix | 713 | 228864 |
| Arabidopsis.Thaliana | 151245 | 66162308 |
| Archaeoglobus | 3757 | 1180343 |
| Bacillus.subtilis | 104992 | 34879326 |
| Caenorhabditis.elegans | 3347 | 2110402 |
| DanioRerio | 24118 | 10557602 |
| Drosophila.melanogaster | 12606 | 8945564 |
| Escherichia.coli | 3983 | 1346730 |
| Helicobacter.pylori | 2392 | 848550 |
| Homo.Sapiens | 140450 | 58477968 |
| Leishmania.major | 8239 | 5285329 |
| M.Xanthus | 5037 | 2043663 |
| Methanosarcina | 2963 | 1039833 |
| MusMusculus | 92857 | 41549270 |
| Myxococcus | 5037 | 2043663 |
| OryzaSativa | 65554 | 25953002 |
| P.Horikoshii | 1583 | 459156 |
| Plasmodiumfalciparum3D7 | 5259 | 4121429 |
| Pyrococcus | 1441 | 462830 |
| Schizosaccharomyces.Pombe | 7711 | 3787977 |
| Staphylococcus.aureus | 1977 | 648703 |
| Streptomyces.coelicolorA3 | 5202 | 1828585 |
| Sulfolobus.solfataricus | 9674 | 2969225 |
| Thermoplasma.acidophilum | 1150 | 379203 |
| ZeaMays | 70650 | 25168719 |

Table 6: Description of the organisms whose complete CDS set has been analyzed in the three reading frames.

We present the coverage ranks within 16 equivalence classes of the set of maximal, self-complementary and $C^3$ circular codes. Within such classes, the best and the worst codes are disjoint sets. In particular, Tables 7 and 8 show the coverage (relative) ranks in frame for the best and worst codes, respectively. Similarly, Tables 9 and 10 show the same analysis where the coding sequences are in the reading frame +1 and the set of codes are obtained from the first circular permutation $\alpha_1(\cdot)$ of the usual set of 216 codes used in frame. Finally, Tables 11 and 12 present the same analysis for frame +2 sequences and the second circular permutation $\alpha_2(\cdot)$ of the set of codes.

| | $X_{173}$ | $X_{23}$ | $X_{98}$ | $X_{166}$ | $X_4$ | $X_{172}$ | $X_{21}$ | $X_{24}$ | $X_{97}$ | $X_{171}$ | $X_3$ | $X_{165}$ | $X_{115}$ | $X_{161}$ | $X_{41}$ | $X_{107}$ |
|---|---|---|---|---|---|---|---|---|---|---|---|---|---|---|---|---|
| AeropyrumPernix | 1 | 1 | 2 | 1 | 1 | 1 | 1 | 1 | 2 | 1 | 1 | 1 | 1 | 1 | 1 | 1 |
| Arabidopsis.Thaliana | 1 | 1 | 1 | 1 | 1 | 1 | 1 | 1 | 1 | 1 | 1 | 1 | 1 | 1 | 1 | 1 |
| Archaeoglobus | 1 | 1 | 2 | 1 | 1 | 1 | 1 | 1 | 2 | 1 | 1 | 1 | 1 | 1 | 1 | 1 |
| Bacillus.subtilis | 1 | 1 | 1 | 1 | 1 | 1 | 1 | 1 | 1 | 1 | 1 | 1 | 1 | 1 | 1 | 2 |
| Caenorhabditis.elegans | 1 | 1 | 1 | 1 | 1 | 1 | 1 | 1 | 1 | 1 | 1 | 1 | 1 | 1 | 2 | 1 |
| DanioRerio | 1 | 1 | 1 | 1 | 1 | 1 | 1 | 1 | 1 | 1 | 1 | 1 | 1 | 1 | 1 | 1 |
| Drosophila.melanogaster | 1 | 1 | 1 | 1 | 1 | 1 | 1 | 1 | 1 | 1 | 1 | 1 | 1 | 1 | 1 | 1 |
| Escherichia.coli | 1 | 1 | 1 | 1 | 1 | 1 | 1 | 1 | 1 | 1 | 1 | 1 | 2 | 1 | 1 | 1 |
| Helicobacter.pylori | 1 | 1 | 1 | 2 | 2 | 1 | 1 | 1 | 1 | 1 | 1 | 1 | 1 | 1 | 2 | 1 |
| Homo.Sapiens | 1 | 1 | 1 | 1 | 1 | 1 | 1 | 1 | 1 | 1 | 1 | 1 | 1 | 1 | 1 | 1 |
| Leishmania.major | 1 | 1 | 2 | 1 | 1 | 1 | 1 | 1 | 1 | 1 | 1 | 1 | 1 | 1 | 1 | 1 |
| M.Xanthus | 1 | 1 | 1 | 1 | 1 | 1 | 1 | 1 | 1 | 1 | 1 | 1 | 1 | 1 | 1 | 1 |
| Methanosarcina | 1 | 1 | 1 | 1 | 1 | 1 | 1 | 1 | 1 | 1 | 1 | 1 | 1 | 1 | 1 | 3 |
| MusMusculus | 1 | 1 | 1 | 1 | 1 | 1 | 1 | 1 | 1 | 1 | 1 | 1 | 1 | 1 | 1 | 1 |
| Myxococcus | 1 | 1 | 1 | 1 | 1 | 1 | 1 | 1 | 1 | 1 | 1 | 1 | 1 | 1 | 1 | 1 |
| OryzaSativa | 1 | 1 | 1 | 1 | 1 | 1 | 1 | 1 | 1 | 1 | 1 | 1 | 1 | 1 | 1 | 1 |
| P.Horikoshii | 1 | 1 | 2 | 1 | 1 | 1 | 1 | 1 | 2 | 1 | 1 | 1 | 1 | 1 | 1 | 1 |
| Plasmodiumfalciparum3D7 | 1 | 1 | 1 | 1 | 1 | 1 | 1 | 1 | 1 | 1 | 2 | 2 | 2 | 2 | 1 | 2 |
| Pyrococcus | 1 | 1 | 2 | 1 | 1 | 1 | 1 | 1 | 2 | 1 | 1 | 1 | 1 | 1 | 1 | 1 |
| Schizosaccharomyces.Pombe | 1 | 1 | 1 | 1 | 1 | 1 | 1 | 1 | 1 | 1 | 1 | 1 | 1 | 1 | 1 | 1 |
| Staphylococcus.aureus | 1 | 1 | 1 | 3 | 1 | 1 | 1 | 1 | 1 | 1 | 2 | 3 | 2 | 2 | 1 | 2 |
| Streptomyces.coelicolorA3 | 1 | 1 | 2 | 1 | 1 | 1 | 1 | 1 | 1 | 1 | 1 | 1 | 1 | 1 | 2 | 3 |
| Sulfolobus.solfataricus | 1 | 1 | 1 | 1 | 1 | 1 | 1 | 1 | 1 | 1 | 1 | 1 | 1 | 1 | 2 | 2 |
| Thermoplasma.acidophilum | 1 | 1 | 2 | 1 | 1 | 1 | 1 | 1 | 2 | 1 | 1 | 1 | 1 | 1 | 1 | 1 |
| ZeaMays | 1 | 1 | 2 | 1 | 1 | 1 | 1 | 1 | 1 | 1 | 1 | 1 | 1 | 1 | 1 | 1 |

Table 7: FRAME 0: relative coverage rank over 25 genomes for the 16 codes identified as best codes. No matter the organism, such codes are almost invariably ranked first within their equivalence class.

| | $X_{192}$ | $X_{87}$ | $X_{53}$ | $X_{191}$ | $X_{86}$ | $X_{195}$ | $X_{91}$ | $X_{57}$ | $X_{54}$ | $X_{208}$ | $X_{90}$ | $X_{194}$ | $X_{136}$ | $X_{207}$ | $X_{93}$ | $X_{146}$ |
|---|---|---|---|---|---|---|---|---|---|---|---|---|---|---|---|---|
| AeropyrumPernix | 8 | 8 | 8 | 8 | 8 | 8 | 8 | 8 | 8 | 8 | 8 | 8 | 8 | 8 | 8 | 8 |
| Arabidopsis.Thaliana | 8 | 8 | 7 | 8 | 8 | 8 | 8 | 8 | 8 | 8 | 8 | 8 | 8 | 8 | 8 | 8 |
| Archaeoglobus | 8 | 8 | 8 | 8 | 8 | 8 | 8 | 8 | 8 | 8 | 8 | 8 | 8 | 8 | 8 | 8 |
| Bacillus.subtilis | 8 | 8 | 7 | 8 | 8 | 8 | 8 | 8 | 8 | 8 | 8 | 8 | 8 | 8 | 8 | 8 |
| Caenorhabditis.elegans | 8 | 8 | 7 | 8 | 8 | 8 | 8 | 8 | 8 | 8 | 8 | 8 | 8 | 8 | 8 | 8 |
| DanioRerio | 8 | 8 | 8 | 8 | 8 | 8 | 8 | 8 | 8 | 8 | 8 | 8 | 8 | 8 | 8 | 7 |
| Drosophila.melanogaster | 8 | 8 | 7 | 8 | 8 | 8 | 8 | 8 | 8 | 8 | 8 | 8 | 8 | 8 | 8 | 8 |
| Escherichia.coli | 8 | 8 | 7 | 8 | 8 | 8 | 8 | 8 | 8 | 8 | 8 | 8 | 8 | 8 | 8 | 7 |
| Helicobacter.pylori | 8 | 8 | 8 | 8 | 8 | 8 | 8 | 8 | 8 | 8 | 8 | 8 | 8 | 8 | 7 | 7 |
| Homo.Sapiens | 8 | 8 | 8 | 8 | 8 | 8 | 8 | 8 | 8 | 8 | 8 | 8 | 8 | 8 | 8 | 8 |
| Leishmania.major | 8 | 8 | 8 | 8 | 8 | 8 | 8 | 8 | 8 | 8 | 8 | 8 | 8 | 8 | 8 | 8 |
| M.Xanthus | 8 | 8 | 8 | 8 | 8 | 8 | 8 | 8 | 8 | 8 | 8 | 8 | 8 | 8 | 8 | 8 |
| Methanosarcina | 8 | 8 | 7 | 8 | 8 | 8 | 8 | 8 | 8 | 8 | 8 | 8 | 8 | 8 | 8 | 8 |
| MusMusculus | 8 | 8 | 8 | 8 | 8 | 8 | 8 | 8 | 8 | 8 | 8 | 8 | 8 | 8 | 8 | 7 |
| Myxococcus | 8 | 8 | 8 | 8 | 8 | 8 | 8 | 8 | 8 | 8 | 8 | 8 | 8 | 8 | 8 | 8 |
| OryzaSativa | 8 | 8 | 8 | 8 | 8 | 8 | 8 | 8 | 8 | 8 | 8 | 8 | 8 | 8 | 8 | 8 |
| P.Horikoshii | 8 | 8 | 8 | 8 | 8 | 8 | 8 | 8 | 8 | 8 | 8 | 8 | 8 | 8 | 8 | 8 |
| Plasmodiumfalciparum3D7 | 7 | 7 | 6 | 8 | 8 | 8 | 8 | 7 | 8 | 7 | 8 | 8 | 8 | 8 | 8 | 8 |
| Pyrococcus | 8 | 8 | 8 | 8 | 8 | 8 | 8 | 8 | 8 | 8 | 8 | 8 | 8 | 8 | 8 | 8 |
| Schizosaccharomyces.Pombe | 8 | 8 | 8 | 8 | 8 | 8 | 8 | 8 | 8 | 8 | 8 | 8 | 8 | 8 | 7 | 8 |
| Staphylococcus.aureus | 7 | 7 | 6 | 8 | 8 | 8 | 7 | 8 | 7 | 7 | 8 | 8 | 8 | 8 | 8 | 8 |
| Streptomyces.coelicolorA3 | 8 | 8 | 8 | 8 | 8 | 8 | 8 | 8 | 8 | 8 | 8 | 8 | 8 | 8 | 8 | 8 |
| Sulfolobus.solfataricus | 8 | 8 | 8 | 8 | 8 | 8 | 8 | 8 | 8 | 8 | 8 | 8 | 8 | 8 | 8 | 8 |
| Thermoplasma.acidophilum | 8 | 8 | 8 | 8 | 8 | 8 | 8 | 8 | 8 | 8 | 8 | 8 | 8 | 8 | 8 | 8 |
| ZeaMays | 8 | 8 | 8 | 8 | 8 | 8 | 8 | 8 | 8 | 8 | 8 | 8 | 8 | 8 | 8 | 8 |

Table 8: FRAME 0: relative coverage rank over 25 genomes for the 16 codes identified as worst codes. No matter the organism, such codes are almost invariably ranked eighth within their equivalence class. They are obtained as the Keto-Amino transformation of the best codes.

| $\alpha_1(\cdot)$ | $X_{173}$ | $X_{23}$ | $X_{98}$ | $X_{166}$ | $X_4$ | $X_{172}$ | $X_{21}$ | $X_{24}$ | $X_{97}$ | $X_{171}$ | $X_3$ | $X_{165}$ | $X_{115}$ | $X_{161}$ | $X_{41}$ | $X_{107}$ |
|---|---|---|---|---|---|---|---|---|---|---|---|---|---|---|---|---|
| AeropyrumPernix | 1 | 1 | 2 | 1 | 1 | 1 | 1 | 1 | 2 | 1 | 1 | 1 | 1 | 1 | 1 | 1 |
| Arabidopsis.Thaliana | 1 | 1 | 2 | 1 | 1 | 1 | 1 | 1 | 2 | 1 | 1 | 1 | 1 | 1 | 2 | 1 |
| Archaeoglobus | 1 | 1 | 3 | 1 | 1 | 1 | 1 | 1 | 2 | 1 | 1 | 1 | 1 | 1 | 2 | 2 |
| Bacillus.subtilis | 1 | 1 | 1 | 1 | 1 | 1 | 1 | 1 | 1 | 1 | 1 | 1 | 1 | 1 | 2 | 2 |
| Caenorhabditis.elegans | 1 | 1 | 2 | 1 | 1 | 1 | 1 | 1 | 2 | 1 | 1 | 1 | 1 | 1 | 1 | 1 |
| DanioRerio | 1 | 1 | 2 | 1 | 1 | 1 | 1 | 1 | 2 | 1 | 1 | 1 | 1 | 1 | 1 | 1 |
| Drosophila.melanogaster | 1 | 1 | 2 | 1 | 1 | 1 | 1 | 1 | 2 | 1 | 1 | 1 | 1 | 1 | 1 | 1 |
| Escherichia.coli | 1 | 1 | 1 | 1 | 1 | 1 | 1 | 1 | 1 | 1 | 1 | 1 | 1 | 1 | 2 | 1 |
| Helicobacter.pylori | 1 | 1 | 1 | 1 | 1 | 1 | 1 | 1 | 1 | 1 | 1 | 1 | 1 | 2 | 3 | 2 |
| Homo.Sapiens | 1 | 1 | 2 | 1 | 1 | 1 | 1 | 1 | 2 | 1 | 1 | 1 | 1 | 1 | 1 | 1 |
| Leishmania.major | 1 | 1 | 2 | 1 | 1 | 1 | 1 | 1 | 2 | 1 | 1 | 1 | 1 | 1 | 1 | 1 |
| M.Xanthus | 2 | 2 | 2 | 1 | 1 | 1 | 1 | 1 | 1 | 1 | 1 | 1 | 1 | 1 | 2 | 2 |
| Methanosarcina | 1 | 1 | 2 | 1 | 1 | 1 | 1 | 1 | 2 | 1 | 1 | 1 | 1 | 1 | 2 | 2 |
| MusMusculus | 1 | 1 | 2 | 1 | 1 | 1 | 1 | 1 | 2 | 1 | 1 | 1 | 1 | 1 | 1 | 1 |
| Myxococcus | 2 | 2 | 2 | 1 | 1 | 1 | 1 | 1 | 1 | 1 | 1 | 1 | 1 | 1 | 2 | 2 |
| OryzaSativa | 1 | 1 | 2 | 1 | 1 | 1 | 1 | 1 | 2 | 1 | 1 | 1 | 1 | 1 | 1 | 1 |
| P.Horikoshii | 1 | 1 | 2 | 1 | 1 | 1 | 1 | 1 | 2 | 1 | 1 | 1 | 1 | 1 | 1 | 2 |
| Plasmodiumfalciparum3D7 | 1 | 1 | 1 | 1 | 1 | 1 | 1 | 1 | 1 | 1 | 2 | 2 | 1 | 1 | 1 | 1 |
| Pyrococcus | 1 | 1 | 2 | 1 | 1 | 1 | 1 | 1 | 2 | 1 | 1 | 1 | 1 | 1 | 1 | 2 |
| Schizosaccharomyces.Pombe | 1 | 1 | 2 | 1 | 1 | 1 | 2 | 1 | 1 | 2 | 1 | 1 | 1 | 1 | 2 | 1 |
| Staphylococcus.aureus | 1 | 1 | 1 | 1 | 1 | 1 | 1 | 1 | 1 | 1 | 1 | 1 | 1 | 1 | 2 | 2 |
| Streptomyces.coelicolorA3 | 2 | 1 | 2 | 1 | 1 | 1 | 1 | 1 | 1 | 1 | 1 | 1 | 1 | 1 | 1 | 3 |
| Sulfolobus.solfataricus | 1 | 1 | 1 | 1 | 1 | 1 | 1 | 1 | 1 | 1 | 1 | 1 | 1 | 1 | 2 | 2 |
| Thermoplasma.acidophilum | 2 | 1 | 3 | 1 | 1 | 1 | 1 | 1 | 2 | 1 | 1 | 1 | 1 | 1 | 1 | 2 |
| ZeaMays | 1 | 1 | 2 | 1 | 1 | 1 | 1 | 1 | 2 | 1 | 1 | 1 | 1 | 1 | 1 | 1 |

Table 9: FRAME +1: relative coverage rank over 25 genomes for the 16 codes identified as best codes. No matter the organism, such codes are almost invariably ranked first within their equivalence class.

| $\alpha_1(\cdot)$ | $X_{192}$ | $X_{87}$ | $X_{53}$ | $X_{191}$ | $X_{86}$ | $X_{195}$ | $X_{91}$ | $X_{57}$ | $X_{54}$ | $X_{208}$ | $X_{90}$ | $X_{194}$ | $X_{136}$ | $X_{207}$ | $X_{93}$ | $X_{146}$ |
|---|---|---|---|---|---|---|---|---|---|---|---|---|---|---|---|---|
| AeropyrumPernix | 8 | 8 | 8 | 8 | 8 | 8 | 8 | 8 | 8 | 8 | 8 | 8 | 8 | 8 | 8 | 8 |
| Arabidopsis.Thaliana | 8 | 8 | 8 | 8 | 8 | 8 | 8 | 8 | 8 | 8 | 8 | 8 | 8 | 8 | 8 | 7 |
| Archaeoglobus | 8 | 8 | 7 | 8 | 8 | 8 | 8 | 8 | 8 | 8 | 8 | 8 | 8 | 8 | 8 | 7 |
| Bacillus.subtilis | 8 | 8 | 8 | 8 | 8 | 8 | 8 | 8 | 8 | 8 | 8 | 8 | 8 | 8 | 8 | 7 |
| Caenorhabditis.elegans | 8 | 8 | 7 | 8 | 8 | 8 | 8 | 8 | 8 | 8 | 8 | 8 | 8 | 8 | 8 | 7 |
| DanioRerio | 8 | 8 | 6 | 8 | 8 | 8 | 8 | 8 | 7 | 8 | 8 | 8 | 8 | 8 | 7 | 8 |
| Drosophila.melanogaster | 8 | 8 | 7 | 8 | 8 | 8 | 8 | 8 | 7 | 8 | 8 | 8 | 8 | 8 | 8 | 7 |
| Escherichia.coli | 8 | 8 | 7 | 8 | 8 | 8 | 8 | 8 | 7 | 8 | 8 | 8 | 8 | 8 | 7 | 7 |
| Helicobacter.pylori | 8 | 8 | 7 | 8 | 8 | 8 | 8 | 8 | 8 | 7 | 8 | 8 | 8 | 8 | 7 | 7 |
| Homo.Sapiens | 8 | 8 | 8 | 8 | 8 | 8 | 8 | 8 | 8 | 8 | 8 | 8 | 8 | 8 | 8 | 8 |
| Leishmania.major | 8 | 8 | 7 | 8 | 8 | 8 | 8 | 8 | 7 | 8 | 8 | 8 | 8 | 8 | 8 | 7 |
| M.Xanthus | 8 | 8 | 6 | 8 | 8 | 8 | 8 | 8 | 7 | 8 | 8 | 8 | 8 | 8 | 6 | 8 |
| Methanosarcina | 8 | 8 | 8 | 8 | 8 | 8 | 8 | 8 | 8 | 8 | 8 | 8 | 8 | 8 | 8 | 8 |
| MusMusculus | 7 | 8 | 6 | 8 | 8 | 8 | 8 | 8 | 7 | 8 | 8 | 8 | 8 | 8 | 7 | 8 |
| Myxococcus | 8 | 8 | 6 | 8 | 8 | 8 | 8 | 8 | 7 | 8 | 8 | 8 | 8 | 8 | 6 | 8 |
| OryzaSativa | 8 | 8 | 8 | 8 | 8 | 8 | 8 | 8 | 8 | 8 | 8 | 8 | 8 | 8 | 8 | 8 |
| P.Horikoshii | 8 | 8 | 8 | 8 | 8 | 8 | 8 | 8 | 8 | 8 | 8 | 8 | 8 | 8 | 8 | 8 |
| Plasmodiumfalciparum3D7 | 7 | 7 | 6 | 8 | 8 | 8 | 8 | 8 | 7 | 8 | 8 | 8 | 8 | 8 | 8 | 8 |
| Pyrococcus | 8 | 8 | 8 | 8 | 8 | 8 | 8 | 8 | 8 | 8 | 8 | 8 | 8 | 8 | 8 | 8 |
| Schizosaccharomyces.Pombe | 8 | 8 | 7 | 8 | 8 | 8 | 8 | 8 | 8 | 8 | 8 | 8 | 8 | 7 | 7 | 7 |
| Staphylococcus.aureus | 7 | 7 | 6 | 8 | 8 | 7 | 7 | 7 | 7 | 8 | 8 | 8 | 8 | 8 | 8 | 8 |
| Streptomyces.coelicolorA3 | 8 | 8 | 8 | 8 | 8 | 8 | 8 | 8 | 8 | 8 | 8 | 8 | 8 | 8 | 7 | 8 |
| Sulfolobus.solfataricus | 8 | 8 | 7 | 8 | 8 | 8 | 8 | 8 | 8 | 8 | 8 | 8 | 8 | 8 | 8 | 8 |
| Thermoplasma.acidophilum | 8 | 8 | 7 | 8 | 8 | 8 | 8 | 8 | 8 | 8 | 8 | 8 | 8 | 8 | 8 | 7 |
| ZeaMays | 8 | 8 | 8 | 8 | 8 | 8 | 8 | 8 | 8 | 8 | 8 | 8 | 8 | 8 | 8 | 8 |

Table 10: FRAME +1: relative coverage rank over 25 genomes for the 16 codes identified as worst codes. No matter the organism, such codes are almost invariably ranked eighth within their equivalence class. They are obtained as the Keto-Amino transformation of the best codes.

| $\alpha_2(\cdot)$ | $X_{173}$ | $X_{23}$ | $X_{98}$ | $X_{166}$ | $X_4$ | $X_{172}$ | $X_{21}$ | $X_{24}$ | $X_{97}$ | $X_{171}$ | $X_3$ | $X_{165}$ | $X_{115}$ | $X_{161}$ | $X_{41}$ | $X_{107}$ |
|---|---|---|---|---|---|---|---|---|---|---|---|---|---|---|---|---|
| AeropyrumPernix | 1 | 1 | 2 | 1 | 1 | 1 | 1 | 1 | 1 | 1 | 1 | 1 | 1 | 1 | 1 | 1 |
| Arabidopsis.Thaliana | 1 | 1 | 2 | 1 | 1 | 1 | 1 | 1 | 2 | 1 | 1 | 1 | 1 | 1 | 1 | 1 |
| Archaeoglobus | 1 | 1 | 2 | 1 | 1 | 1 | 1 | 1 | 2 | 1 | 1 | 1 | 1 | 1 | 1 | 1 |
| Bacillus.subtilis | 1 | 1 | 1 | 1 | 1 | 1 | 1 | 1 | 1 | 1 | 1 | 1 | 1 | 1 | 2 | 1 |
| Caenorhabditis.elegans | 1 | 1 | 2 | 1 | 1 | 1 | 1 | 1 | 2 | 1 | 1 | 1 | 1 | 1 | 1 | 1 |
| DanioRerio | 1 | 1 | 2 | 1 | 1 | 1 | 1 | 1 | 2 | 1 | 1 | 1 | 1 | 1 | 1 | 1 |
| Drosophila.melanogaster | 1 | 1 | 3 | 1 | 1 | 1 | 1 | 1 | 2 | 1 | 1 | 1 | 1 | 1 | 1 | 1 |
| Escherichia.coli | 1 | 1 | 1 | 1 | 1 | 1 | 1 | 1 | 1 | 1 | 1 | 1 | 1 | 1 | 1 | 1 |
| Helicobacter.pylori | 1 | 1 | 1 | 1 | 1 | 1 | 1 | 1 | 1 | 1 | 1 | 1 | 1 | 2 | 2 | 2 |
| Homo.Sapiens | 1 | 1 | 2 | 1 | 1 | 1 | 1 | 1 | 2 | 1 | 1 | 1 | 1 | 1 | 1 | 1 |
| Leishmania.major | 1 | 1 | 2 | 1 | 1 | 1 | 1 | 1 | 2 | 1 | 1 | 1 | 1 | 1 | 1 | 1 |
| M.Xanthus | 2 | 1 | 2 | 1 | 1 | 1 | 1 | 1 | 2 | 1 | 1 | 1 | 1 | 1 | 1 | 1 |
| Methanosarcina | 1 | 1 | 2 | 1 | 1 | 1 | 1 | 1 | 1 | 1 | 1 | 1 | 1 | 1 | 1 | 1 |
| MusMusculus | 1 | 1 | 2 | 1 | 1 | 1 | 1 | 1 | 2 | 1 | 1 | 1 | 1 | 1 | 1 | 1 |
| Myxococcus | 2 | 1 | 2 | 1 | 1 | 1 | 1 | 1 | 2 | 1 | 1 | 1 | 1 | 1 | 1 | 1 |
| OryzaSativa | 1 | 1 | 2 | 1 | 1 | 1 | 1 | 1 | 2 | 1 | 1 | 1 | 1 | 1 | 1 | 1 |
| P.Horikoshii | 1 | 1 | 1 | 1 | 1 | 1 | 1 | 1 | 1 | 1 | 1 | 1 | 1 | 1 | 1 | 1 |
| Plasmodiumfalciparum3D7 | 1 | 1 | 1 | 1 | 1 | 1 | 1 | 1 | 1 | 1 | 2 | 2 | 1 | 1 | 2 | 1 |
| Pyrococcus | 1 | 1 | 1 | 1 | 1 | 1 | 1 | 1 | 1 | 1 | 1 | 1 | 1 | 1 | 1 | 1 |
| Schizosaccharomyces.Pombe | 1 | 1 | 1 | 1 | 1 | 1 | 2 | 1 | 1 | 1 | 2 | 1 | 1 | 1 | 2 | 1 |
| Staphylococcus.aureus | 1 | 1 | 1 | 1 | 1 | 1 | 1 | 1 | 1 | 1 | 2 | 2 | 2 | 2 | 2 | 2 |
| Streptomyces.coelicolorA3 | 2 | 1 | 2 | 1 | 1 | 2 | 1 | 1 | 2 | 1 | 1 | 1 | 1 | 1 | 1 | 3 |
| Sulfolobus.solfataricus | 1 | 1 | 1 | 1 | 1 | 1 | 1 | 1 | 1 | 1 | 1 | 1 | 1 | 1 | 1 | 2 |
| Thermoplasma.acidophilum | 2 | 1 | 2 | 1 | 1 | 1 | 1 | 1 | 2 | 1 | 1 | 1 | 1 | 1 | 1 | 1 |
| ZeaMays | 1 | 1 | 2 | 1 | 1 | 1 | 1 | 1 | 2 | 1 | 1 | 1 | 1 | 1 | 1 | 1 |

Table 11: FRAME +2: relative coverage rank over 25 genomes for the 16 codes identified as best codes. No matter the organism, such codes are almost invariably ranked first within their equivalence class.

| $\alpha_2(\cdot)$ | $X_{192}$ | $X_{87}$ | $X_{53}$ | $X_{191}$ | $X_{86}$ | $X_{195}$ | $X_{91}$ | $X_{57}$ | $X_{54}$ | $X_{208}$ | $X_{90}$ | $X_{194}$ | $X_{136}$ | $X_{207}$ | $X_{93}$ | $X_{146}$ |
|---|---|---|---|---|---|---|---|---|---|---|---|---|---|---|---|---|
| AeropyrumPernix | 8 | 8 | 8 | 8 | 8 | 8 | 8 | 8 | 8 | 8 | 8 | 8 | 8 | 8 | 8 | 8 |
| Arabidopsis.Thaliana | 8 | 8 | 7 | 8 | 8 | 8 | 8 | 8 | 7 | 8 | 8 | 8 | 8 | 8 | 7 | 7 |
| Archaeoglobus | 8 | 8 | 8 | 8 | 8 | 8 | 8 | 8 | 8 | 8 | 8 | 8 | 8 | 8 | 8 | 7 |
| Bacillus.subtilis | 8 | 8 | 8 | 8 | 8 | 8 | 8 | 8 | 8 | 8 | 8 | 8 | 8 | 8 | 8 | 7 |
| Caenorhabditis.elegans | 8 | 8 | 7 | 8 | 8 | 8 | 8 | 8 | 7 | 8 | 8 | 8 | 8 | 8 | 8 | 8 |
| DanioRerio | 8 | 8 | 7 | 8 | 8 | 8 | 8 | 8 | 7 | 8 | 8 | 8 | 8 | 8 | 8 | 8 |
| Drosophila.melanogaster | 7 | 8 | 7 | 8 | 8 | 8 | 8 | 8 | 7 | 8 | 8 | 8 | 8 | 8 | 8 | 7 |
| Escherichia.coli | 8 | 8 | 7 | 8 | 8 | 8 | 8 | 8 | 7 | 8 | 8 | 8 | 8 | 8 | 8 | 6 |
| Helicobacter.pylori | 8 | 8 | 7 | 8 | 8 | 8 | 8 | 8 | 7 | 7 | 8 | 8 | 8 | 8 | 8 | 8 |
| Homo.Sapiens | 8 | 8 | 7 | 8 | 8 | 8 | 8 | 8 | 8 | 8 | 8 | 8 | 8 | 8 | 8 | 8 |
| Leishmania.major | 8 | 8 | 7 | 8 | 8 | 8 | 8 | 8 | 7 | 8 | 8 | 8 | 8 | 8 | 8 | 6 |
| M.Xanthus | 8 | 8 | 6 | 8 | 8 | 8 | 8 | 8 | 7 | 8 | 8 | 8 | 8 | 8 | 8 | 8 |
| Methanosarcina | 8 | 8 | 7 | 8 | 8 | 8 | 8 | 8 | 7 | 8 | 8 | 8 | 8 | 8 | 8 | 8 |
| MusMusculus | 7 | 8 | 6 | 8 | 8 | 7 | 8 | 7 | 7 | 8 | 8 | 8 | 8 | 8 | 7 | 8 |
| Myxococcus | 8 | 8 | 6 | 8 | 8 | 8 | 8 | 8 | 7 | 8 | 8 | 8 | 8 | 8 | 8 | 8 |
| OryzaSativa | 8 | 8 | 7 | 8 | 8 | 8 | 8 | 8 | 8 | 8 | 8 | 8 | 8 | 8 | 8 | 8 |
| P.Horikoshii | 8 | 8 | 7 | 8 | 8 | 8 | 8 | 8 | 8 | 8 | 8 | 8 | 8 | 8 | 8 | 8 |
| Plasmodiumfalciparum3D7 | 7 | 7 | 6 | 8 | 8 | 8 | 8 | 8 | 7 | 8 | 8 | 8 | 8 | 8 | 8 | 8 |
| Pyrococcus | 8 | 8 | 8 | 8 | 8 | 8 | 8 | 8 | 8 | 8 | 8 | 8 | 8 | 8 | 8 | 8 |
| Schizosaccharomyces.Pombe | 8 | 8 | 7 | 8 | 8 | 8 | 8 | 8 | 7 | 8 | 8 | 8 | 8 | 8 | 7 | 6 |
| Staphylococcus.aureus | 7 | 7 | 6 | 8 | 8 | 7 | 7 | 7 | 6 | 7 | 8 | 8 | 8 | 8 | 8 | 8 |
| Streptomyces.coelicolorA3 | 8 | 8 | 8 | 8 | 8 | 8 | 8 | 8 | 8 | 8 | 8 | 8 | 8 | 8 | 8 | 8 |
| Sulfolobus.solfataricus | 8 | 8 | 7 | 8 | 8 | 8 | 8 | 8 | 8 | 8 | 8 | 8 | 8 | 8 | 8 | 8 |
| Thermoplasma.acidophilum | 8 | 8 | 8 | 8 | 8 | 8 | 8 | 8 | 8 | 8 | 8 | 8 | 8 | 8 | 8 | 7 |
| ZeaMays | 8 | 8 | 7 | 8 | 8 | 8 | 8 | 8 | 8 | 8 | 8 | 8 | 8 | 8 | 8 | 8 |

Table 12: FRAME +2: relative coverage rank over 25 genomes for the 16 codes identified as worst codes. No matter the organism, such codes are almost invariably ranked eighth within their equivalence class. They are obtained as the Keto-Amino transformation of the best codes.

*2.3. The universal properties of circular codes are absent in introns.*

The structure uncovered in coding sequences is completely absent in introns as it is shown in Table S13, where we present the mean coverage over 225 intron sequences of A.thaliana. Clearly, there is no organization implied by circular codes within introns.

| **coverage** | $X_{173}$ | $X_{176}$ | $X_{203}$ | $X_{206}$ | $X_{183}$ | $X_{182}$ | $X_{193}$ | $X_{192}$ |
|---|---|---|---|---|---|---|---|---|
| frame 0 | 29.5 | 28.9 | 30.3 | 29.7 | 29.4 | 29.3 | 29.6 | 29.6 |
| frame +1 | 29.1 | 29.5 | 28.8 | 29.2 | 29.2 | 29.3 | 28.9 | 29.0 |
| frame +2 | 28.9 | 29.3 | 28.9 | 29.3 | 29.0 | 29.1 | 29.2 | 29.2 |
| **relative rank** | $X_{173}$ | $X_{176}$ | $X_{203}$ | $X_{206}$ | $X_{183}$ | $X_{182}$ | $X_{193}$ | $X_{192}$ |
| frame 0 | 4 | 1 | 8 | 7 | 3 | 2 | 5 | 5 |
| frame +1 | 4 | 8 | 1 | 5 | 5 | 7 | 2 | 3 |
| frame +2 | 1 | 7 | 1 | 7 | 3 | 4 | 5 | 5 |

Table 13: Mean coverage (upper panel) and relative rank (lower panel) of the 8 circular codes forming the equivalence class presented in Table S2 computed over 225 intron sequences of A. thaliana.

In conclusion, each circular code has a distinct degree of coverage with respect to the species-specific codon usage of distinct organisms, according also to the GC content. Notably, however, there are recurring properties, linking the coverage inside equivalence classes with the set of chemical transformations of the codons of the codes.

*2.5. Circular codes and codon influence on protein expression*

Indeed, there is no evident correlation between single codon influence and single codon usage (Figure 2).

*2.6. Circular code motifs are absent in the mRNA 5'-head and 3'-tail sequences*

If circular code motifs/properties have a role in translation, then a differential coverage of the codons belonging to circular codes could apply as a function of position in the coding sequence. In Figure 3 we plotted the coverage of codes $X_{173}$ (blue solid line) and $X_{192}$ (red solid line) over rolling windows of 5 codons, computed over the first 100 codons of each complete coding sequence of the 25 organisms described in Table S6. Remarkably, both for code $X_{173}$ and $X_{192}$ there is a transient initial span of around 40 codon positions after which the rolling coverage over 5 codons reaches the value of the global coverage over the entire genome and fluctuates around it. While for code $X_{173}$ the rolling coverage for the first positions is always lower than the global coverage, the rolling coverage for code $X_{192}$ starts at a higher level with respect to the global coverage and decreases towards it. This appears to be a universal feature shared by all the organisms. The same is true for rolling windows up to 30 codons with no significant differences. The effect of the total codon content in the tail of the sequence was also reported to be influential (Boël et al., 2016) on expression.

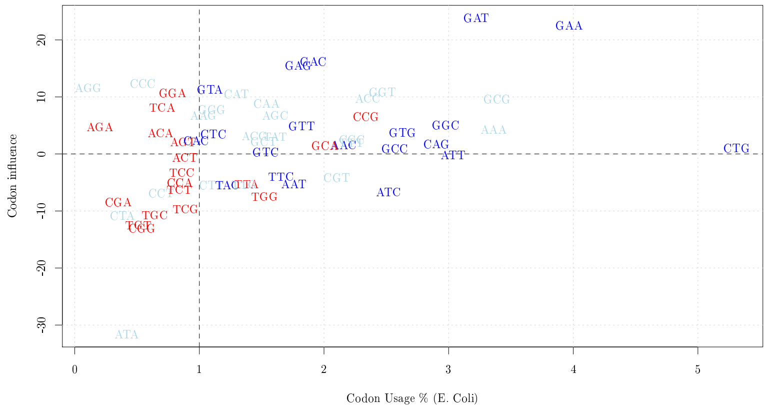

Figure 2: Codon usage in E. coli (percent) versus codon influence. The codons coloured in blue and in red belong to circular codes $X_{173}$ and $X_{192}$, respectively. They are the best and worst codes within the set of 8 codes forming the equivalence class shown in Table S3.

Accordingly, we also observed a universal tail effect in the coverage of coding sequences (Figure 4).

##### *2.7. circular code properties correlate with the S/W character of the first two nucleotides of the codon*

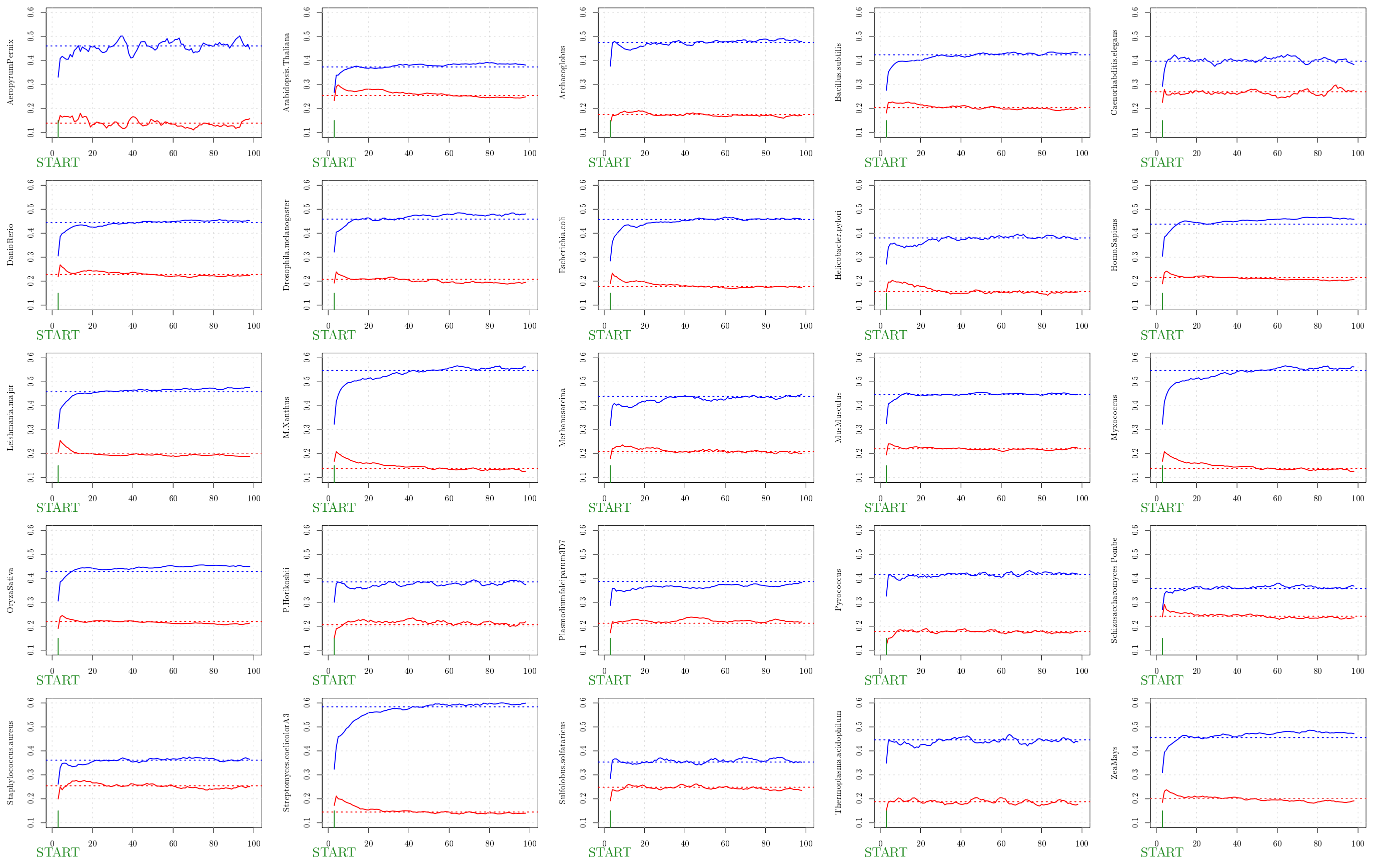

Figure 3: Rolling coverage computed on the first 100 codon positions, averaged over the whole set of complete coding sequences of 25 genomes. The blue and red solid lines correspond to code $X_{173}$ and $X_{192}$, respectively. The dotted lines correspond to the global coverage of the codes over the whole genome.

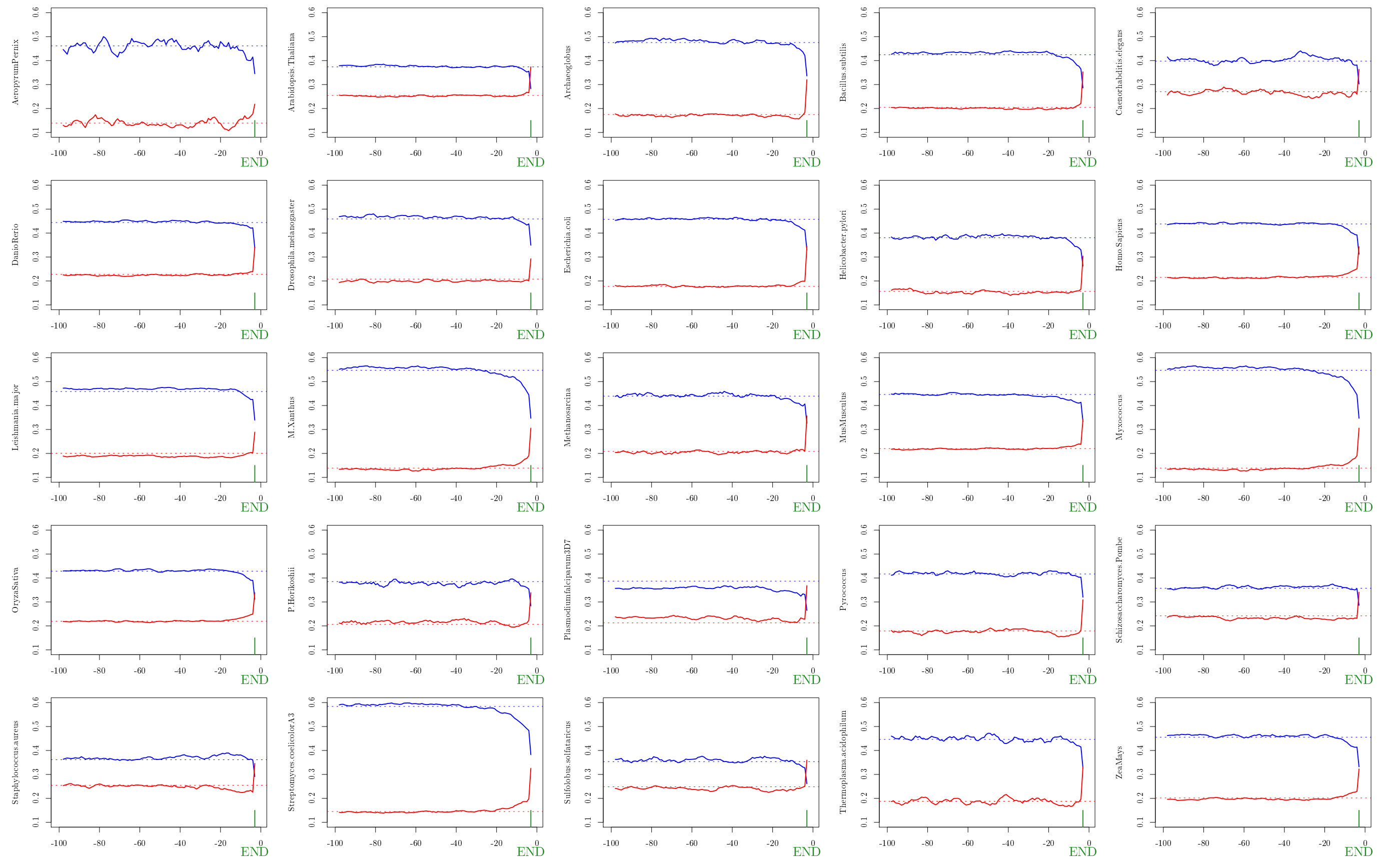

Figure 4: Rolling coverage computed on the last 100 codon positions, averaged over the whole set of complete coding sequences of 25 genomes. The blue and red solid lines correspond to code $X_{173}$ and $X_{192}$, respectively. The dotted lines correspond to the global coverage of the codes over the whole genome.

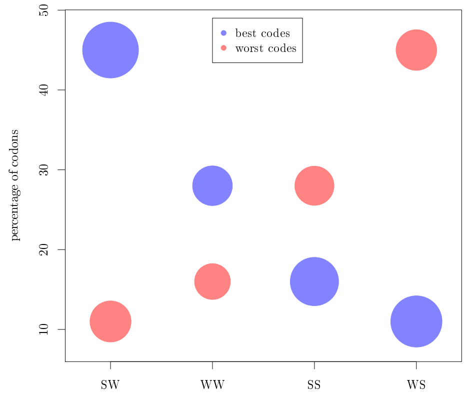

Figure 5: Comparison of codon composition of best codes (blue) and worst codes (red) according to the S/W chemical dichotomy of the first two nucleotides of the codon. The area of the bubbles is proportional to the average codon influence.

Michel, C., Pirillo, G., & Pirillo, M. (2008). A relation between trinucleotide comma-free codes and trinucleotide circular codes. *Theoretical Computer Science*, *401*, 17 – 26.

Nakamura, Y., Gojobori, T., & Ikemura, T. (1997). Codon usage tabulated from the international DNA sequence databases. *Nucleic Acids Research*, *25*, 244. doi:10.1093/nar/25.1.244.

Nirenberg, M., & Matthaei, J. (1961). The dependence of cell-free protein synthesis in E. coli upon naturally occurring or synthetic polyribonucleotides. *Proceedings of the National Academy of Sciences*, *47*, 1588–1602.
